# ACENet: a graph neural network for predicting enzyme pHmin using structure and sequence insights

**DOI:** 10.64898/2026.03.11.711222

**Authors:** Shantong Hu, Tong Pan, Jiayu Wang, Huan Yee Koh, Zhikang Wang, Yumeng Zhang, Lingxuan Zhu, Zhuoqian Li, Geoffrey I. Webb, Jiangning Song, Guimin Zhang

## Abstract

The acid activity of enzymes, characterized by the minimum pH at which enzymes remain active (pHmin), is crucial for industrial applications in acidic environments. However, the rational design of acid-active enzymes remains challenging due to limited understanding of sequence-structure- activity relationships under acidic conditions. Here, we propose ACENet, a graph neural network that predicts enzyme pHmin by integrating surface features of protein structures with evolutionary representations derived from the large-scale protein language model ESM-2. ACENet achieved a Pearson correlation coefficient of 0.85 on the test dataset, significantly outperforming other deep learning baseline models and maintains stable pHmin predictions under various conditions. Even on a subset of the dataset with less than 20% homology, the PCC remains above 0.5, with an RMSE (Root mean square error) less than 1.4. ACENet also present excellent performance in the annotation of pHmin for homologous proteins and the predictive screening of minimal active pH in protein mutants. Remarkably, ACENet could identify the catalytic region as key determinants of acid activity through residue-level interpretability analysis. Overall, ACENet accelerates the development of highly efficient biocatalysts for diverse applications where acidic conditions predominate.

## 1. Introduction

Many enzymes need to adapt to a wide pH range to function under practical or physiological conditions. The activity and stability of enzymes under acidic conditions have long been a focus of attention in some industrial enzyme applications. For instance, the enzymes fed to livestock and poultry need to be able to pass through the entire digestive tract and remain highly active in an acidic to neutral pH range, or be able to tolerate the low pH of gastric juice and function under neutral pH in the intestines [1]. Another example is the corn steep liquor (CSL) deep-processing industry, where the pH of CSL is below 4.0. This process requires enzymes with high activity under acidic conditions, such as amylase and protease [2]. In some enzymatic reactions, such as those catalyzed by deacetylase, acetic acid is continuously produced to lower the reaction pH [3]. This requires the engineered deacetylase to maintain high activity and stability during the process to maintain a high substrate-product conversion rate. Therefore, enzymes with higher acid activity and stability are crucial to improving the catalytic efficiency in these applications.

Mining enzymes directly from microorganisms that thrive in acidic environments is a common strategy for obtaining enzymes with robust activity under acidic conditions. However, this approach has its own limitations, as the target enzyme may not be present in these microorganisms; even if it is present, the enzyme may not exhibit high activity or stability under acidic conditions because the pH values of the intracellular environment is often maintained at a neutral level[4]. Another strategy is to optimize existing enzymes through protein engineering, but the physicochemical principles governing enzyme function at extreme pH conditions remain complex and poorly understood. Current empirical efforts typically fall into two categories; one involves rationally designing the catalytic pocket residues and adjusting their pKa values to shift the enzyme’s pH activity profile. The other method is to lower the enzyme’s surface charges by introducing more acidic residues (aspartic acid (D) or glutamic acid (E)). For instance, Wang et al. used PROPKA to design the A270K/N271H mutant, which reduced the pKa of the catalytic residue D174 from 8.05 to 3.68, thereby reducing the optimal pH of Amy7C by 2.0 units[5], and Yan et al. used the Rosetta Supercharge algorithms to modify the glucose oxidase of *Aspergillus niger* and reduced its pHopt (optimum pH) by 1.0 (from 6.0 to 5.0) via the Q241E/R499E double mutations[6]. Besides, other studies also showed that mutations introduced into catalytical pockets often reduces enzyme activity, and changes in pHopt do not necessarily improve acid stability. For example, Xie et al. engineered a pullulanase double mutant (Q13H/I25E) whose pHopt shifted from 6.4 to 5.0, but the mutant completely lost its activity at pH 4.0[7].

Recent advances in deep learning have led to transformative applications across a wide range of biological tasks, including chemical space exploration[8], gene expression analysis[9], and prediction of parameters such as enzyme affinity and turnover rate[10]. Among these advances, graph neural networks (GNNs) have shown great promise in protein-related research. Notably, models such as ProteinMPNN[11], LigandMPNN[12], and AlphaFold[13] have used GNNs as core components for learning protein features, making significant progress in *de novo* protein design and structure prediction. In addition, protein language models (PLMs) such as ESM-2 have also introduced a new paradigm for protein sequence modeling[14]. These large-scale Transformer- based models are trained on billions of protein sequences and structure-function data, and can directly extract protein evolution and functional information from protein sequences[15]. Recent studies have shown that PLMs can predict protein stability[16], enzymatic function[17], and mutation effects[18], making them as a promising tool for enzyme engineering and functional optimization[15]. Several deep learning tools, such as Ophpred[19], Seq2pHopt[20], and EpHod[21], have been developed to predict the optimal pH of proteins. Nevertheless, as mentioned above, changing the pHopt value alone does not necessarily improve the acid activity and stability of enzymes, particularly at pH values below 4. This underscores the need for a dedicated deep learning framework capable of systematically predicting the acid activity and stability of enzymes under highly acidic conditions.

To address these challenges, we introduced the “pHmin” metric, inspired by the pH range annotations in the BRENDA database, defined as the lowest pH value at which an enzyme maintains detectable activity. Lower pHmin values indicate greater acid tolerance, reflecting higher activity and stability under acidic conditions. To facilitate systematic research, we compiled a new benchmark dataset and developed ACENet, a deep learning method based on BRENDA annotations. ACENet uses protein structure data as input and integrates sequence-based evolutionary features and surface structural information to predict pHmin values, thereby determining the acid activity and stability of the protein. The model was validated with test datasets from various sources, demonstrating strong predictive performance and robustness in predicting enzyme pHmin values. These results offer promise for mining and engineering enzymes with enhanced acid activity and stability.

## 2. Results

### 2.1 Overview of ACENet framework

To systematically predict enzyme stability and activity under acidic conditions, we constructed a comprehensive benchmark dataset comprising 20,284 enzyme samples, along with their associated sequences, structures, EC numbers, experimental pH-activity profiles, and annotations. This dataset was randomly split into training and testing sets at a 9:1 ratio, and a five-fold cross-validation was applied to the training set to validate the model (**Fig. 1A**; see the Methods section for details).

**Figure 1.**
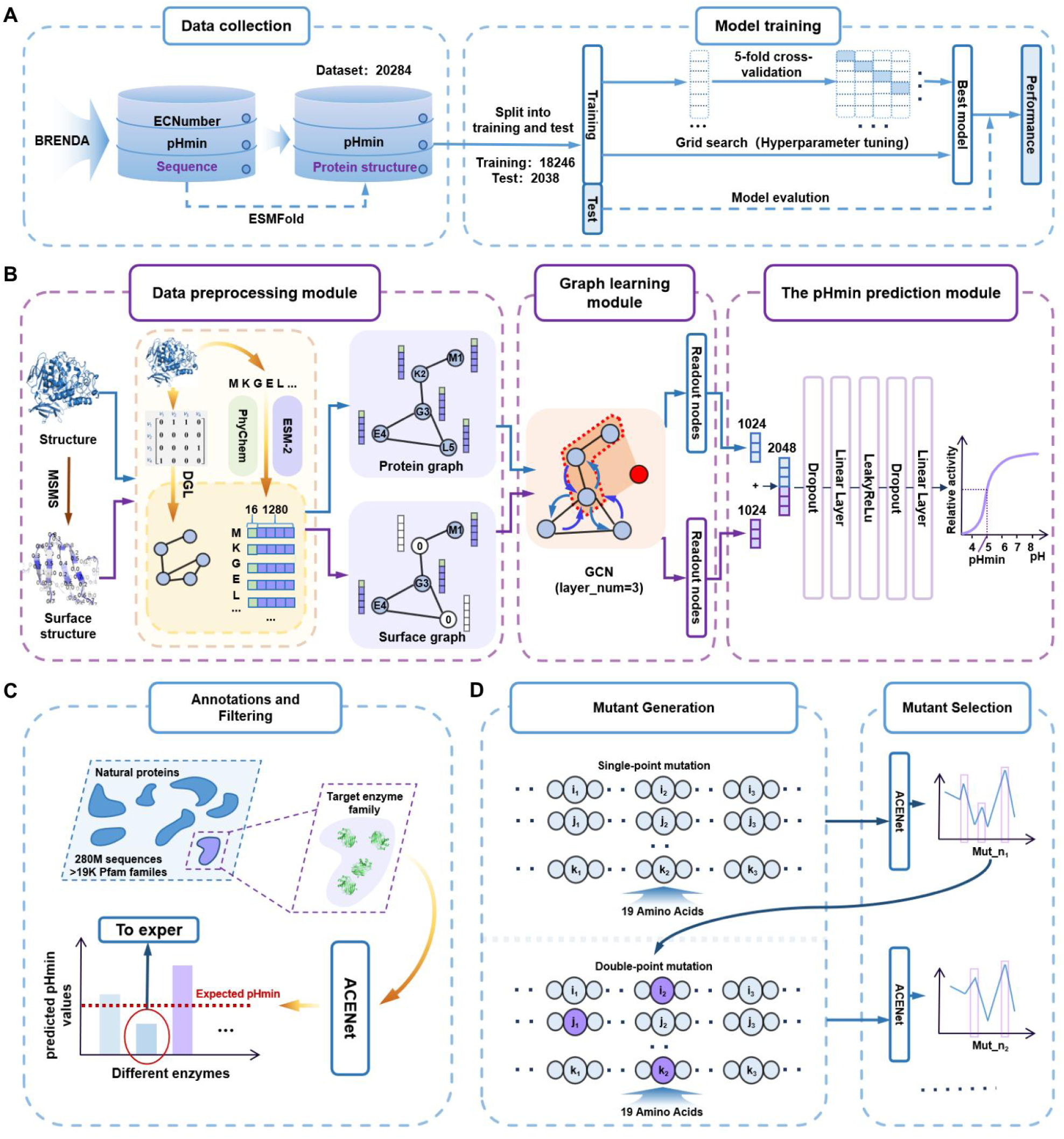
Overview of the ACENet framework for protein acid stability prediction. (A)Data collection and model training. Protein sequences, structures, and pHmin values were collected from the BRENDA database, and protein structures were predicted using ESMFold. This dataset contains 20,284 samples, which were divided into a training dataset (18,246) and a test dataset (2,038). Model training includes a 5-fold cross-validation strategy, hyper-parameter optimization via grid search, and performance evaluation to determine the optimal prediction model. **(B) ACENet architecture.** The framework consists of three modules: (i) a data preprocessing module, which uses DGL and ESM-2 embeddings to process protein sequence and structural information into graph representations. A protein backbone graph and surface graphs are constructed to capture structural features. (ii) a graph learning module, in which a three-layer graph convolutional network (GCN) extracts relevant features from the protein and surface graphs and enhances the representation with residue-level nodes. (iii) a pHmin prediction module, which uses a multi-layer perceptron (MLP) to predict the minimum pH value at which a protein remains stable. **(C) Overview of the protein annotation and screening process.** Target protein families can be mined from nature and annotated using ACENet to screen for samples that maintain activity under specified acidic pH conditions for experimental validation, thereby reducing costs. **(D) Systematic mutant generation and screening framework.** We developed an ACENet-based mutant design and screening framework that automatically performs iterative saturation mutagenesis and rounds of screening from candidate mutation sites to obtain ideal mutants.

The architecture of ACENet is shown in **Fig. 1B**, which comprises three main components: a data preprocessing module, a graph learning module, and a pHmin prediction module. During the preprocessing phase, the 3D structure of the protein is converted into a graph representation. For each protein, a two-layer graph was constructed: in the main graph, nodes represent individual amino acids with sequence-derived features, and edges represent spatial proximity between residues. Simultaneously, a surface graph was generated by focusing on solvent-accessible residues and their local neighborhoods to capture surface-specific features relevant to acid activity and stability. The feature vector of each node is enriched by combining evolutionary embeddings and a joint representation of physicochemical properties. Specifically, we combined sequence embeddings from the ESM-2 transformer model [15], which provides an evolutionary informed residue profile, as well as descriptors of their chemical properties and structural context. To enrich the model’s evolutionary context, we introduced ESM-2 embeddings to leverage information learned from large-scale protein sequence databases. The protein graph and surface-based graph were processed in parallel by a graph learning module consisting of a multi-layer graph convolutional network (GCN) to propagate and aggregate node-level information. The representations obtained from the two graph paths were then concatenated and fed into a feed-forward prediction module (comprising a fully connected layer with dropout regularization) to generate the final prediction.

To demonstrate the versatility of ACENet for pHmin prediction, we conducted extensive validation to support its application in two key areas: large-scale enzyme pHmin annotation (**Fig. 1C**) and rational mutant design and screening (**Fig. 1D**). The annotation workflow (**Fig. 1C**) systematically identifies target enzymes with acid tolerance using a large-scale protein databases containing 280 million sequences from over 19 K Pfam families[22]. The mutant design framework (**Fig. 1D**) implements a systematic approach where ACENet identifies better mutants, followed by comprehensive iterative site-saturation mutagenesis. Mutants were then prioritized based on their predicted pHmin, streamlining candidate selection for experimental validation.

### 2.2 Performance Evaluation of ACENet

To assess the model’s fit to the training data, we monitored its performance across the training epochs. As shown in **Fig. 2A**, the Spearman correlation coefficient (SPCC) for pHmin predictions consistently improved with increasing training epochs. At epoch 35, the Root Mean Square Error (RMSE), SPCC, and Pearson correlation coefficient (PCC) values of 0.83 ± 0.04, 0.79 ± 0.01, and 0.80 ± 0.02 (**Tab. S1**) on the validation set for the nine models trained with 5-fold cross-validation are 0.83 ± 0.03, 0.79 ± 0.01, and 0.80 ± 0.01, respectively (**Tab. S2**). The model achieved nearly identical performance on both datasets, demonstrating its prediction stability on the benchmark dataset. Moreover, a high degree of agreement was observed between the predicted pHmin values and the experimentally determined pHmin values.

**Figure 2.**
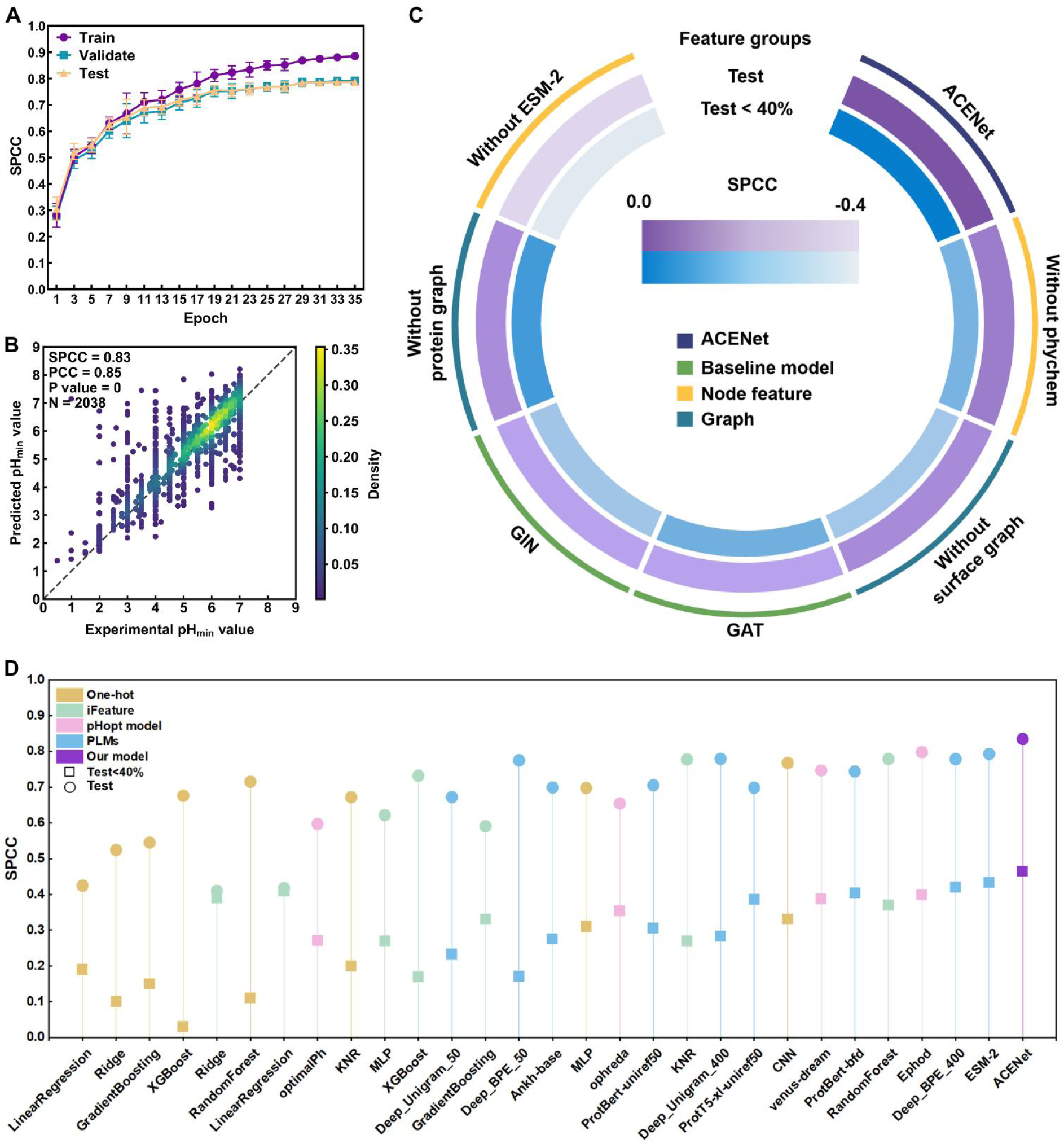
Evaluation of ACENet’s performance for pHmin prediction. (A) Training performance at different epochs. ACENet’s performance was evaluated during training, validation, and testing. The SPCC is plotted against the number of training epochs; error bars represent the mean and standard deviation of 5-fold cross- validation. The model exhibites steady performance improvement and convergence. **(B) Comparison of predicted and experimental pHmin values.** The predicted values for the test dataset were compared with the experimentally measured pHmin values. The density of the data points is indicated by color intensity, with darker colors indicating higher density. **(C) Ablation study of ACENet.** By systematically removing key features (including protein graphs, surface graphs, physicochemical properties (PhyChem), and ESM-2 embeddings), the impact of different components on model performance was analyzed. The purple and blue ring represent the performance degradation after removing each component on the full test set and test set (test < 40%), respectively. The baseline model replaces the core ACENet with alternative graph neural network architectures, such as the Graph Attention Network (GAT) and the Graph Isomorphism Network (GIN). Model performance is measured using the SPCC metrics, where lighter colors indicate greater importance of the feature. **(D) Performance comparison of the best model from each method.** ACENet was benchmarked against regression models (including XGBoost, RandomForest, GradientBoosting, Ridge, KNR, LinearRegression, and MLP, CNNs), several PLMs from Venus Factory and prior methods— EpHod, OphPred, Venus-Dream, opHReda. “Test” and “Test < 40% “ represent the complete test set and test set sequences and structures with an identity of less than 40% to the training set, respectively.

The final ACENet model was trained on the full training dataset and evaluated on an independent test set. Comparison of the predicted pHmin values with the experimental pHmin values (**Fig. 2B**) showed SPCC, PCC, and RMSE values of 0.84, 0.85, and 0.72 (**Tab. S3**) respectively, all exceeding the results of five-fold cross-validation. These findings confirm a strong positive correlation between model predictions and experimental data, further supporting the reliability and robustness of the model. In addition, we conducted ablation experiments to systematically evaluate the function of individual model components in the ACENet model (**Fig. 2C**) on full testing set and testing set sequences and structures with less than 40% identity (test < 40%) to the training set. Notably, removing ESM-2 features from ACENet resulted in a substantial drop in performance, underscoring their critical contribution to the model (**Table S3**).

Furthermore, we compared ACENet with several representative machine-learning methods, deep- learning models and prior methods— EpHod[21], OphPred[19], Venus-Dream[23], opHReda[24] using the same experiment setup. On these benchmarks, ACENet achieved the highest performance compared to other approaches on both the full test set and test set (test <40%) to the training set, however, SPCC only scores 0.47 on the testing set (test < 40%), slightly better than the recently published EpHod (0.40) and Venus-Dream (0.39) (**Fig. 2D**). This assessment convincingly demonstrates the superior predictive power of ACENet for enzyme pHmin (**Table S4**). Comparative analysis also showed that ESM-2, a transformer-based protein language model, provides the most informative sequence representation. Significantly, ACENet’s superior performance remains even when other models adopt the same feature representations, further highlighting the inherent effectiveness of this architecture in addressing this specific bioinformatics challenge.

### 2.3 Evaluation of predictive generalization and stability

Next, we conducted a comprehensive evaluation of ACENet on the entire testing dataset to rigorously assess its generalizability, especially under the following challenging conditions: (i) low sequence identity between the testing and training datasets; (ii) enzymes with different EC Numbers; (iii) targets requiring significantly lower pHmin values (pH < 5). Given the inherent sparsity of available data, models developed in this setting often show larger prediction errors when applied to data that deviates significantly from the training distribution. As expected, the results show that model performance gradual declined with decreasing sequence identity or structural similarity (**Fig. 3A**). For the most divergent sub-datasets (sequence identity < 0.2 or TM-score < 0.2), SPCC values fall below the ideal value (**Fig. 3A**), while PCC and RMSE values remain above 0.50 and below 1.32(**Tab. S5-S6**), respectively. These findings suggest that ACENet maintains good predictive accuracy even for proteins that are phylogenetically distant from the training dataset.

**Figure 3.**
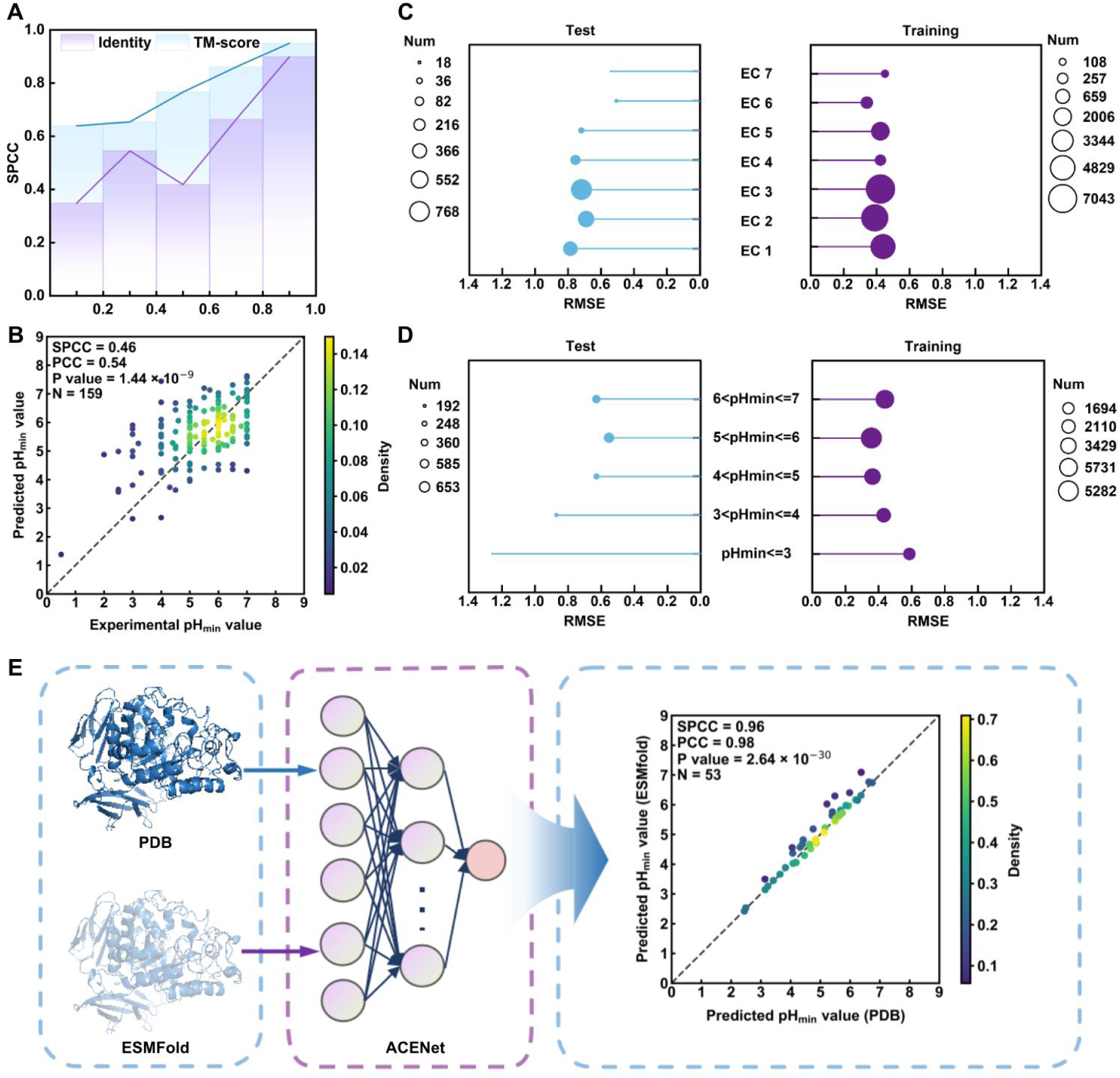
Evaluation of ACENet’s generalization and prediction stability. (A) The impact of sequence and structural similarity on model performance. ACENet’s generalization was assessed by examining its performance on the test dataset at varying levels of sequence identity and structural similarity (TM-score). The shaded region represents the confidence intervals. **(B) ACENet’s performance on low-similarity proteins.** ACENet was tested on a sub-dataset of proteins with sequence identity and structural similarity below 40%. The scatter plots compare the predicted values with the experimental pHmin values, with color intensity representing data density. (**C**) **Prediction accuracy across different EC Numbers.** 1: oxidoreductases, 2: transferases, 3: hydrolases, 4: lyases, 5: isomerases, 6: ligases, 7: translocases. **(D) Prediction accuracy across different pH stability ranges.** Model performance was evaluated by classifying proteins by acid stability, with pHmin thresholds ranging from < 3 to 6 ≤ pHmin < 7. RMSE values for the training and test data are plotted, with dot size representing the number of proteins in each category. (E) **Effect of experimentally solved structures and predicted structures on pHmin prediction**.

To further validate the robustness of ACENet, an independent testing sub-dataset was generated using 159 samples with both sequence identity and TM score below 0.4. The model showed a significant positive correlation between the predicted pHmin values and the experimentally measured values (PCC=0.54; SPCC=0.46; **Fig. 3B**). Notably, predictions for pHmin between 5-7 are more reliable and more consistent with the true values, likely due to the larger sample size within this interval. Conversely, the predictions for pHmin values less than 4 have larger errors and show a systematic tendency to overestimate.

Furthermore, we divided the test and training sets according to 7 main EC numbers and used ACENet to predict the pHmin of enzymes. The results showed that the model had good prediction accuracy for enzymes from different families on the training set, with an RMSE value of approximately 0.4 (Table S7). For the test set, the RMSE values for EC 6 and EC 7 ranged from 0.4 to 0.6, while the RMSE values for enzymes from EC 1 to EC 5 families ranged from 0.6 to 0.8 (Figure 3C, Table S7). Such low RMSE values demonstrate that ACENet can effectively learn pHmin-related features across different enzyme families.

In the original dataset, the class distribution was obviously uneven, with the number of samples decreasing gradually in the lower pHmin range (Fig S1). For instance, there are 5,282 samples in the pHmin range of 6-7. while there are only 1,694 training samples with pHmin less than 3 (**Tab. S8)**. To quantify the effect of this uneven distribution, a hierarchical error analysis was conducted by calculating the RMSE of the pHmin intervals in the training and test datasets. In the training set, the RMSE showed a moderate dependence on the sample size. Results indicated that ACENet performed regions with high data density (pHmin 4-7) compared to regions with sparse pHmin value distribution (pHmin≤4) (**Fig. 3D, Tab. S8**). The errors in the test dataset reflected the limitations of the training data distribution. The RMSE is significantly lower within the pHmin range of 4 to 7. In contrast, when pHmin≤3, the RMSE increased by 44.3% compared to that of pHmin≤4 in the test dataset (**Tab. S9**), indicating that the model has relatively low prediction accuracy for data with pHmin≤3.

Beyond evaluating generalizability on existing data, we also investigated the potential impact of differences in input structural accuracy on prediction outcomes. Considering that a substantial portion of the PDB database has been incorporated into the ESMFold training set, we tested the model’s performance on a subset of CASP competition data using both experimentally solved PDB structures and ESMFold-predicted structures as inputs. The root mean square deviation (RMSD) values between the predicted structures and experimental structures of 53 collected proteins were also compared, with an average value of 6.61(**Fig. S2C).** The pHmin prediction results showed a remarkably high correlation between predictions from both input types. Intriguingly, we observed a systematic trend: predictions based on ESMFold yield higher pHmin values compared to those based on experimental structures (**Fig. 3E**). Despite the current limited validation dataset, this overestimation trend may be beneficial for screening applications: when applying the pHmin threshold for data filtering, the higher predicted values of the ESMFold structures may help reduce the false positive rates. These analysis suggest that the increase in prediction errors may be caused by the uneven distribution of the dataset and inaccurate structural, while the ACENet architecture can maintain basic prediction fidelity even in underrepresented categories, thereby enhancing the model’s generalization ability under data skew conditions of real-world prediction

### 2.4 ACENet can predict pHmin for secreted and non-secreted microbial proteins

To further validate the reliability of the model predictions, we collected complete protein data from 4 strains of archaea, 11 gram-positive bacteria, and 4 Gram-negative bacteria, totaling of 70,151 protein sequences, and calculated their host pH growth range and optimal growth range (detailed information for each strain is provided in **Table S10**). All proteins were classified as secreted or non-secreted categories. We used the ACENet developed in this study and the recently released optimal pH prediction tool EpHod[25] to predict the pHmin and pHopt values of these proteins, respectively. All prediction results were compiled by strain.

Statistical analysis (**Fig. 4**) revealed that the pHmin and pHopt distributions for secreted proteins are lower than those for non-secreted proteins. This pattern is particularly pronounced in the archaea. With the exception of *Ignicoccus islandicus*, which has a relatively high growth pH range (3.8-6.5), all tested archaea can grow below pH 2.0, with an optimal growth pH around 3.0. This observation may be attributed to the fact that archaea predominantly thrive in extremely acidic conditions. Additionally, the pHmin values predicted by ACENet for proteins from all strains were notably lower than the predicted pHopt (**Fig. 4**), consistent with biological reality and indirectly supporting the rationality of the model’s predictions by comparing the predicted distribution ranges. Notably, the predicted pHmin values for secreted proteins from each strain consistently fell within the strain’s growth pH range, with the majority falling below the strain’s optimal growth pH.

**Figure 4.**
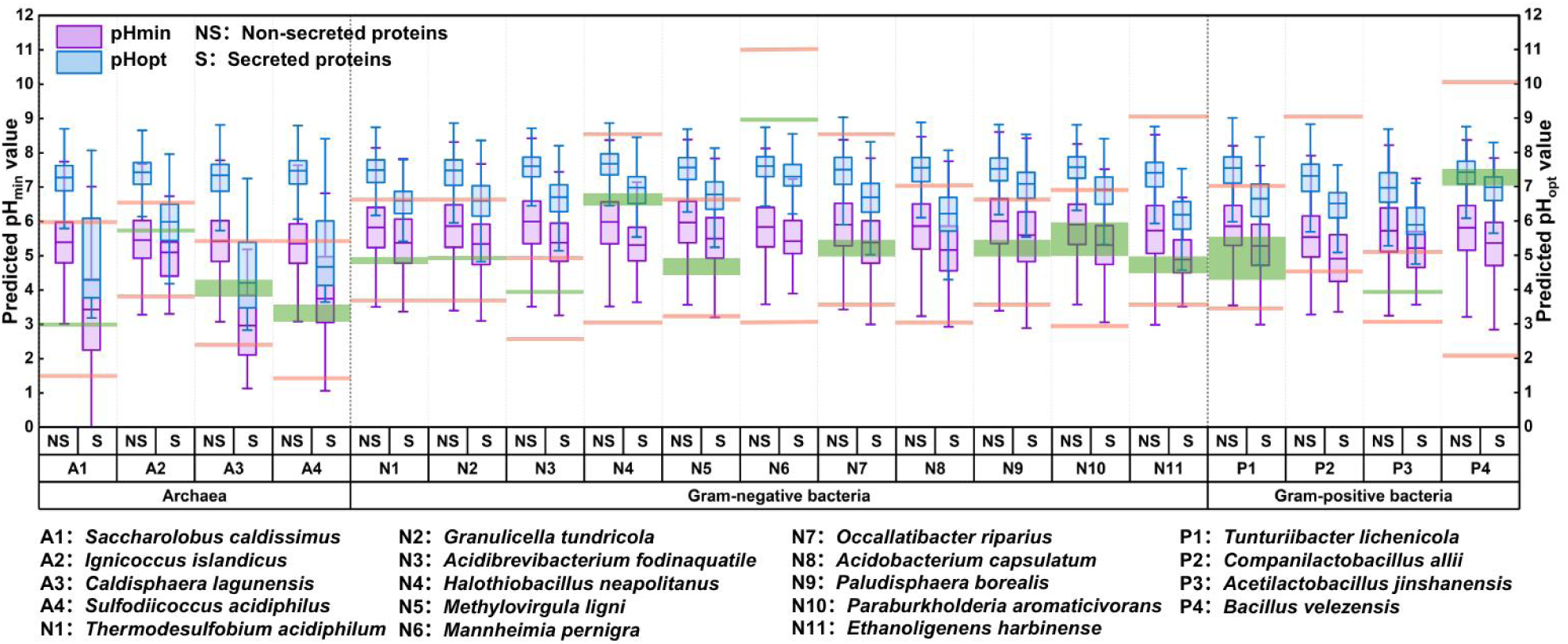
Systematic evaluation of the pH activity profiles across complete proteomes of different strains capable of growing in acid environments. The distribution of predicted pHmin and pHopt values for the complete proteomes of each strain is shown. The purple box plots show the distributions of predicted pHmin value, and the blue box plots showf the distribution of predicted pHopt values for all proteins. The orange upper and lower horizontal boundaries represent the pH growth range for each strain, whereas the light green elements (boxes or lines) indicate the optimal pH for bacterial growth. I denotes intracellular proteins and S denotes secreted proteins.

### 2.5 ACENet-based enzyme pHmin annotation and interpretability analysis

ACENet was then applied to two specific enzymes-glutathione transferase and carbonic anhydrase- for further study. Glutathione transferase is the only known enzyme capable of opening the epoxide ring of vomitoxin[26], but loses its activity under acidic conditions [27]. Carbonic anhydrase plays a crucial role in carbon neutralization efforts to combat global warming by converting CO_2_ into carbonate and bicarbonate ions, improving their bioavailability and conversion rate [28]. The acid resistance and activity of these enzymes are critical for the degradation of vomitoxin and CO_2_ utilization, respectively. We used ACENet to comprehensively annotate the pHmin values of these two families and systematically compared the predicted values with experimental data (**Fig. 5A-B**). This analysis revealed a strong correlation between the predicted and experimental results, with SPCC exceeding 0.70 and PCC exceeding 0.79 for both families. However, when pHmin ≤ 3, the predicted values showed a systematic overestimation compared to the experimental values. This discrepancy likely stemmed from the relatively small number of samples with pHmin values below 4 compared to those above 4 in our dataset.

**Figure 5.**
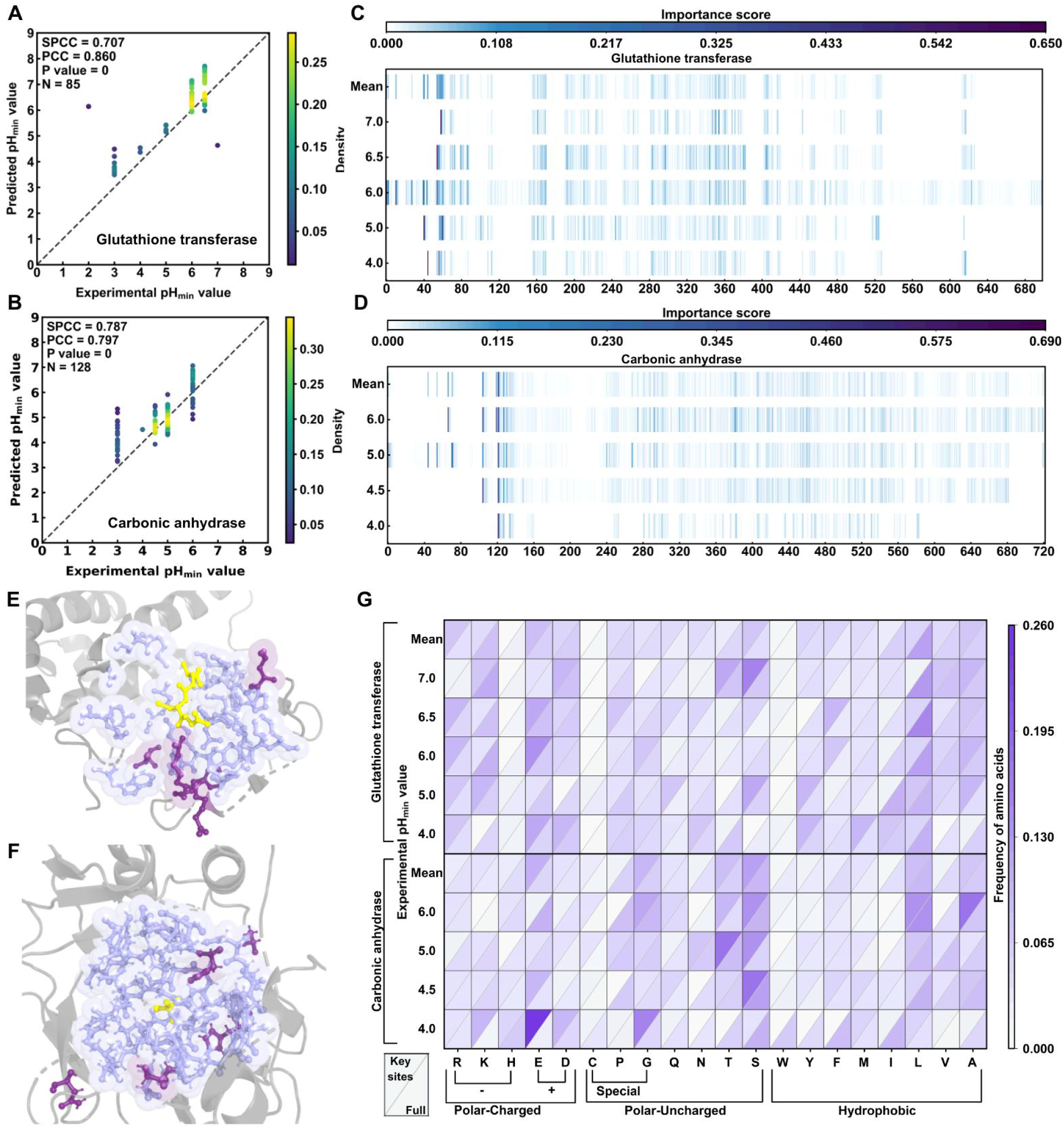
Deep learning-based prediction and interpretation of pHmin for different enzyme families. (A-B) Prediction accuracy for specific enzyme families. Scatter plots show the correlation between predicted pHmin and experimental values for (A) glutathione transferase (N=85) and (B) carbonic anhydrase (N=128). Color intensity reflects the density of data points, with darker colors indicating higher frequency of values. **(C-D) Residue importance analysis of glutathione transferase and carbonic anhydrase.** Residue importance scores were used to identify key amino acid positions that contribute to pHmin predictions. Heatmaps show the average importance scores across all proteins in each enzyme class for (C) glutathione transferase and (D) carbonic anhydrase averaged. Residue positions with higher importance scores indicate regions where perturbations significantly affect pHmin predictions. Analysis was performed by aligning multiple sequences within each enzyme family and normalizing for sequence lengths to maintain consistency. **(E-F) Structural mapping of ACENet interpretability results for glutathione transferase and carbonic anhydrase.** Yellow color indicates the ligand. The blue sphere region represents the catalytic pocket (within 8 Å of the substrate binding site). Purple sticks indicate the top 5 key residues identified by ACENet interpretability analysis. Residues in purple spheres are located within the catalytic pocket. **(G) Amino acid composition of key sites at different pHmin groups.** The amino acid frequency of key functional sites at different pHmin groups within an enzyme family are compared. Key sites were identified with the top 50 positions with the highest importance scores. Heatmaps show the amino acid composition of key sites (upper triangle) and the overall sequence composition (lower triangle). Amino acids are grouped according to their chemical properties, including polar charged residues, polar uncharged residues, special residues, and hydrophobic residues. Darker colors indicate higher frequencies of occurrence.

Next, we conducted a residue-level interpretability analysis of ACENet. To ensure the reliability of the subsequent interpretability analysis, we excluded residue importance calculations for pH ≤ 3 range, allowing for clearer identification of systematic patterns in regions with more reliable predictions. The residue importance calculations for the model identified by ACENet showed that regions with high importance scores were mainly localized at or near the N-terminus. Specifically, the key amino acids for glutathione transferase are primarily located at positions 40-80 (**Fig. 5C**), whereas the critical residues for carbonic anhydrase are concentrated between positions 120-140 (**Fig. 5D**). To visualize the distribution of important residues in the structure, we retrieved structural data and functional annotations for these two enzymes from RCSB PDB. Remarkably, the top 5 amino acids, ranked by importance scores, are mainly distributed within the enzyme’s active pocket (within 8 Å of the substrate binding site) and on the surrounding surface (**Fig. 5E-F**). This finding demonstrates that, without any prior knowledge or guidance about catalytic sites, the model unexpectedly demonstrated, through detailed studies of these enzyme families and a comprehensive literature review, that the key residues identified in both classes of enzymes are well localized within their respective catalytic and binding domains. This spontaneous localization to functionally critical regions significantly enhances the credibility of the model and provides strong confidence in its application to mutant prediction tasks.

Remarkably, a comparative analysis of the amino acid composition of these key residues with that of the complete protein sequences revealed significant compositional differences across the pHmin range (**Fig. 5G**). Specifically, enzymes with lower pHmin values exhibited a higher proportion of acidic amino acids, aspartic acid (D) and glutamic acid (E), at key residue positions (**Tab. S11**), compared to the complete sequence (**Tab. S12**). This finding is consistent with the previously reported higher frequencies of acidic amino acids in acid-resistant enzymes[29]. Additionally, enzymes with lower pHmin values displayed a significantly lower frequency of serine (S) at key residue positions (**Tab. S11**), suggesting a previously unreported compositional preference.

Consistent observations were also obtained when we extended the interpretability analysis to the entire dataset, comparing the amino acid compositions of key residues and the complete protein sequences across all proteins (**Tab. S13-14**). Notably, ACENet was trained without any annotations of the enzyme’s catalytic pockets, demonstrating that the model autonomously learns to focus on amino acids close to the catalytic region, resulting in higher predictive accuracy.

### 2.6 ACENet for pHmin prediction of enzyme mutants

We further investigated the ability of ACENet to quantify the effects of amino acid substitutions on the pHmin of individual enzymes. During the benchmark dataset collection process, we identified a large number of amino acid substitution data, which were individually screened and analyzed. A curated dataset containing mutation data for 10,490 protein samples was divided into wild-type and mutant enzyme groups. The model’s ability to achieve good fit on these datasets represents the first step in validating its application to mutant prediction tasks. Statistical analysis revealed that ACENet achieved PCC and SPCC values exceeding 0.90, with statistically significant P values for both the wild-type (**Fig. 6A**) and mutant protein datasets (**Fig. 6B**) (P < 0.001). Furthermore, the absolute prediction errors of pHmin values on the mutant dataset exhibited a tight distribution centered around zero (**Fig. 6C**), indicating robust prediction reliability and providing strong confidence in the predicted pHmin values of mutant proteins.

**Figure 6.**
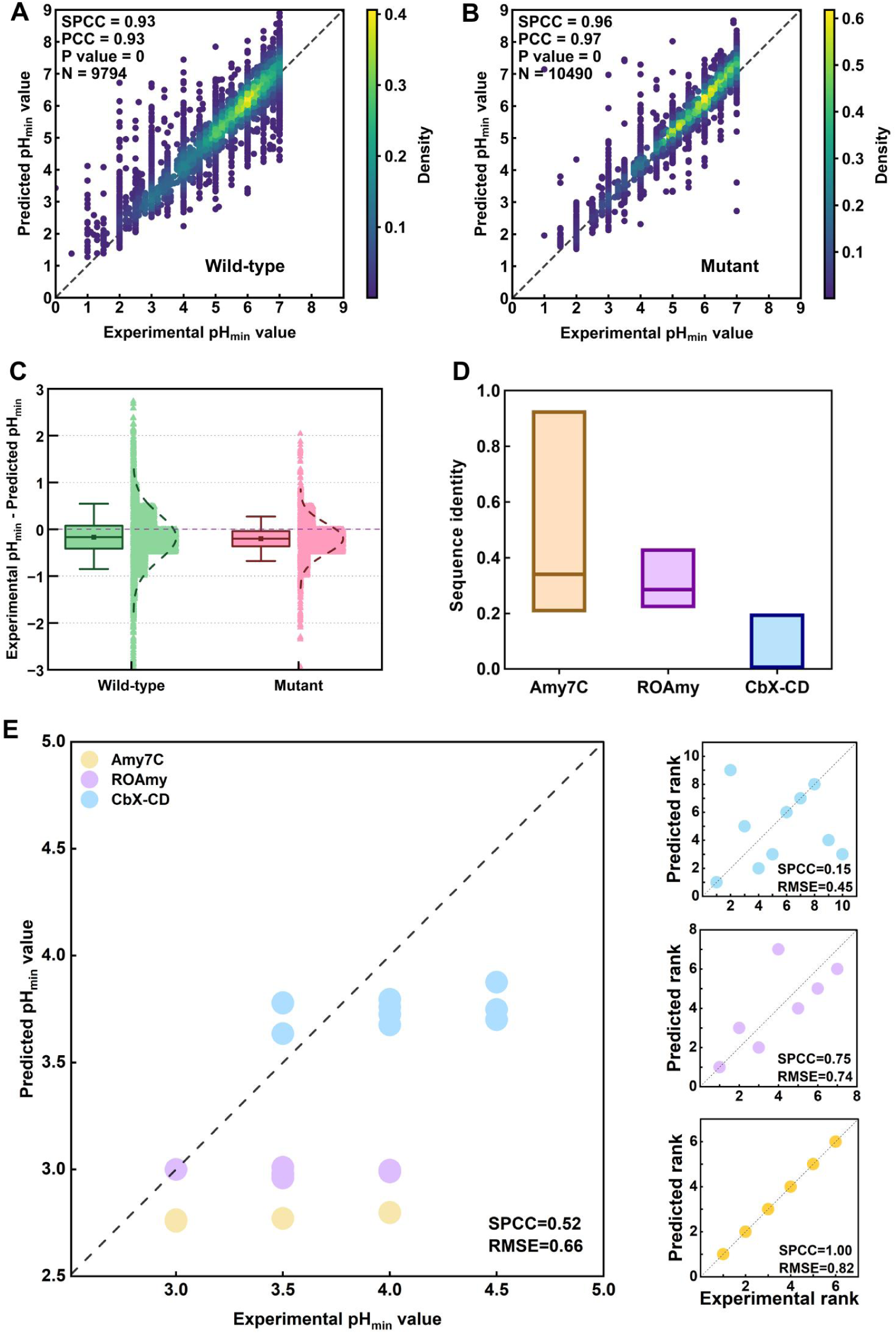
Prediction of pHmin for wild-type and mutant enzymes using ACENet. (A-B) Prediction performance for wild-type and mutant enzymes. Scatter plots compare the predicted and experimental pHmin values for (A) wild-type enzymes (N=9,794) and (B) mutant enzymes (N=14,090). Color intensity represents the density of data points, with darker colors indicating higher density. **(C) Bias analysis of predicted pHmin values.** Box plots show the distribution of differences between the experimental and predicted pHmin values for wild-type and mutant enzymes. **(D) Sequence alignment results of the three collected wild-type enzymes with the training dataset.** The y-axis represents sequence homology, and the box plot range indicates the distribution of sequence similarity when aligning each enzyme template with the entire training dataset using MMseqs2. **(E) Comparative analysis of predicted and experimental results for three enzymes and their mutants.** Yellow, purple, and blue dots represent Amy7C, ROAmy, and CbX-CD and their respective mutants. To provide a detailed visualization of the model’s prediction performance based on relative activity curve position, we derived a detailed relative ranking of pHmin values from the curves and compared this ranking with the predicted values.

We also evaluated the correlation between the predicted values and experimental values of 1,056 mutants in the test set (Fig. S3A), with an SPCC of 0.94. For these data, the minimum sequence similarity with the training set was 0.63, and the sequence identities of most data were higher than 0.8. Moreover, we divided the mutant data in the test set into two parts according to whether they appeared in the training set. The results showed that the presence of templates in the training set significantly affected the model’s prediction accuracy, with the RMSE difference of approximately 0.4 (Fig. S3B). These results imply that the presence of templates in the training set significantly reduces the prediction error of ACENet.

We then selected three glycoside hydrolases (GHs) from different families not present in the training dataset: Amy7C (GH13), ROAmy (GH31), and CbX-CD (GH46), which were selected due to their high acid dependence in industrial applications. Mutant studies of each enzyme were documented in the literature, including their relative activities at different pH values. A total of 23 data points were collected and complied, including Amy7C and its 5 mutants, ROAmy and its 6 mutants, and CbX-CD and its 9 mutants (**Tab. S15**). Sequence alignment analysis revealed that Amy7C had a maximum homology of 0.92 with proteins in the training dataset, but most homologous proteins clustered around 0.32. ROAmy had a peak homology of 0.44 with proteins in the training set, with homologous proteins predominantly distributed around 0.27(**Fig. 6D**). Note that the minimum computational threshold for MMseqs2 is 0.2, and smaller values cannot calculated; therefore, the values for CbX-CD were interpolated, as no proteins with homology greater than 0.2 were found in the CbX-CD training dataset (**Fig. 6D**). This means that CbX-CD does not share detectable homology with any proteins in the training dataset.

These data were then used for prediction by ACENet, revealing three trends (**Fig. 6E**): (1) The more relevant data in the training set, the stronger the model’s ranking ability. (2) The higher the actual pHmin, the smaller the model’s prediction error. (3) The higher the homology between the target protein and the proteins in the training set, the higher the prediction accuracy. These findings are consistent with previous research results and existing report[30]. Admittedly, collecting mutant relative activity curve data from the literature is challenging, resulting in a relatively small test dataset that may affect the reliability of the model’s predictions. Researchers with experimental data for their target enzymes can use the scripts available on GitHub to add this data to the training set to improve ACENet’s prediction performance for their target enzymes.

## 3. DISCUSSION

The acid stability and catalytic efficiency of enzymes under acidic conditions remain a major bottleneck in industrial biotechnology and agricultural applications[31]. In particular, in feed enzyme production, gastric pH compatibility is crucial[32]. To address the limitation, we developed ACENet (Acid Catalysis Enzyme Network), a novel deep learning framework that can directly predict the pHmin values of enzymes. ACENet fills a critical gap in enzyme engineering: the lack of reliable computational tools for predicting enzyme acid tolerance. The success of this model stems from three key innovations that distinguish it from existing approaches. First, by using pHmin rather than pHopt as the target metric, we directly address industrial needs, where sustained activity under harsh conditions is more important than peak performance. Second, the integration of graph neural networks and protein language models enables the captures of local structural constraints and evolutionary conserved patterns that govern acid stability. Third, a dual-graph architecture separately models bulk properties and surface interactions of proteins, reflecting the distinct roles of structural integrity and solvent accessibility in pH tolerance. We also systematically compiled pH activity profiles from comprehensive enzyme databases, creating a curated dataset covering diverse enzyme families and pH ranges.

ACENet’s superior performance across diverse enzyme families suggests that it captures the complex hierarchical relationships governing acid stability, as confirmed by our ablation studies, which show that each feature category contributes significantly to prediction accuracy. We acknowledge that the available training data represents only a small fraction of the natural protein sequence space and is significantly biased toward mesophilic organisms and neutral pH. This sampling bias reflects the ecological distribution of characterized organisms—acidophiles are a minority compared to neutrophiles— resulting in a database dominated by enzymes with limited acid tolerance. To alleviate this concern, we conducted a comprehensive evaluation across sequence- based segmentation, structural similarity variations, and different pHmin ranges to validate the broad applicability of the model. ACENet maintained consistent performance (RMSE < 1.3) across all evaluation subsets, including those representing extreme pH ranges or unusual sequence compositions (**Fig. 3C**) The model was able to discriminate differences in pHmin between highly similar proteins, highlighting its sensitivity to subtle structural determinants of acid stability. To further challenge the model’s generalization capabilities, we collected a comprehensive proteome dataset of acidophilic microorganisms and leveraged principles from microbial evolution to identify enzyme families that are naturally acid-adapted. We further annotated these proteomes for secretion signals, predicted pHmin values, and pH optima using multiple computational approaches (**Fig. 4**). Strain-level analysis of the predicted distributions provided compelling evidence for the robustness of the model, with ACENet, similar to EpHod [25], accurately capturing known trends in acidophilic adaptation while identifying novel candidates for experimental characterization.

In interpretability analyses, in addition to the expected focus on solvent-exposed residues that directly interact with the acidic environment, ACENet consistently highlighted residues within the catalytic and binding sites across diverse enzyme families. This observation is consistent with recent findings that ESM-2 embeddings implicitly encode functional information[33], suggesting that our graph neural network architecture can successfully extract and use catalytic signatures for acid stability prediction. Our analysis also confirms established protein engineering strategies, particularly highlighting the stabilizing role of surface acidic residues (E/D) in low-pHmin enzymes, transforming ACENet from a predictive tool to a mechanistic guide for rational enzyme design. These interpretable predictions provide valuable hypotheses for understanding the molecular determinants of pH-dependent enzyme function.

Improving the acid activity of target enzymes has been a major concern of research, as optimizing existing, high-catalytic activity enzymes is much more efficient than extensive library screening. We experimentally demonstrate that ACENet performs better in predicting mutants with homologous templates in the training set. This result provides a reference for ACENet in screening sequence saturation mutations for enzyme acid stability and activity. We also provided a script on GitHub to help researchers add their experimental data to the training set to optimize ACENet’s predictions for target enzymes.

Despite these significant advances, several important limitations remain. Data quality remains a persistent challenge—redundancies, annotation errors, and measurement inconsistencies in the source databases will be propagated through model training[34, 35]. Furthermore, observed discrepancies between predicted and experimental pHmin shifts for point mutations may partially reflect intrinsic variability in pH-activity assays rather than pure model limitations. We systematically re-analyzed the BRENDA database entries and found substantial heterogeneity in experimental conditions, buffer systems, and substrate choices, which are factors that can significantly influence reported pHmin values. Future directions should prioritize the following key areas: expanding the collection of high-quality experimental datasets covering extreme pH ranges, incorporating substrate-specific effects and cofactor dependencies, and accurately modelling pH- induced conformational transitions. Combined with molecular dynamics simulations, this can provide a deeper understanding of the pH-dependent stability mechanisms at the atomic level[36]. In addition, transfer learning approaches can further improve the prediction accuracy of underrepresented enzyme families by leveraging the growing corpus of protein stability data.

In summary, ACENet represents a significant advancement in computational enzyme engineering, bridging the gap between sequence, structure, and function to accurately predict enzyme acid tolerance. The model’s interpretable architecture provides both a practical tool for enzyme discovery and theoretical insights into the mechanisms of adaptation. As deep learning methods continue to mature and experimental datasets expand, ACENet and similar approaches will play increasingly important role in developing acid-tolerant biocatalysts for sustainable industrial processes.

## 4. Experimental section

### 4.1 Benchmark dataset

The benchmark dataset was retrieved from the BRENDA Enzyme Database on July 10, 2024, using a custom script via an application programming interface (API). This generated a comprehensive dataset containing organism information, EC number, protein identifier (UniProt ID), enzyme type, and pHmin value. Protein sequences queries were conducted using two methods: for entries with UniProt ID information, amino acid sequences were retrieved using Biopython v.1.78 (https://biopython.org/) via the UniProt API. For entries without UniProt ID, amino acid sequences were retrieved from both UniProt and the BRENDA databases based on their EC number and organism information. Specifically, pHmin, protein sequences, and EC numbers were extracted from the dataset, and enzyme entries with pHmin values below 7 were retained. Subsequently, the protein sequences were input into ESMFold[15] to generate protein structure information. Both sequence and structure information are used for network development and testing.

### 4.2 ACENet Framework

#### 4.2.1 Data preprocessing module

Each protein PDB file is processed into graph-structure data using the pipeline provided by the Eggplanck team (winner of the Novozymes Enzyme Stability Prediction competition on Kaggle https://www.kaggle.com/code/gyozzza/nesp-gnn-data). Specifically, the α-carbon atom coordinates of each residue were first extracted, and pairwise Euclidean distances were calculated. Atoms were then classified as connected or disconnected using a preset 8Å threshold. The graph structure was initialized using the DGL platform based on the binary adjacency matrix. For node features, we used a combined encoding strategy based on evolutionary information and physicochemical properties. The evolutionary information was derived from a pre-trained large-scale language model ESM-2, representing each amino acid as a 1280-dimensional vector. For physicochemical features, our encoding strategy integrates attributes such as dissociation constant, charge, solvent accessibility, molecular weight, and specific atomic ratios of amino acid groups; representing each amino acid as a 16-dimensional vector. Finally, these vectors were concatenated to generate a 1296-dimensional amino acid-level node feature.

In addition to the above pretreatment, this study also introduced protein surface information. It is worth noting that the protein surface refers to the part of the protein’s three-dimensional structure that is exposed to the solvent (typically water), and is the primary interface for its interacting with the solvents, other molecules, or ligands in the environment. To characterize the protein surface, we used the MSMS[37] and MaSIF[38] workflows to calculate the solvent accessibility of each amino acid residue, and thus identified surface-exposed residues. Next, a surface map was generated by zeroing the eigenvectors of nodes (residues not on the surface) based on the previously constructed graph derived from the complete protein structure. This resulted in a map containing only information about surface-exposed amino acid residues.

The complete protein map and the surface-specific map were then processed in parallel in subsequent modules for feature learning and prediction.

#### 4.2.2 Graph learning module

The graph learning module consists of a three-layer neural network built upon the Graph Convolutional Network (GCN) [39]. GCN learns node representations by aggregating information from adjacent nodes in the graph. The message passing process of GCN can be formulated as follows:

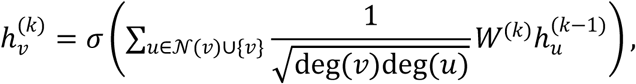

where 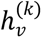 is the feature representation of node at layer 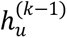 is the feature representation of node (a neighbor of or itself) at layer − *k* − 1 *N(v)* represents the set of adjacent nodes of *v* deg(*v*) and deg(*u*) denote the degrees of nodes and, respectively. *W*^(k)^ is the learnable weight matrix for layer *k*, and σ is a nonlinear activation function, such as LeakyReLU.

Following the GCNs, a multiple layer perceptron (MLP) consisting of fully connected layers and nonlinear activation functions was applied for feature recalibration. Finally, a graph-level representation was obtained by aggregating node features and converting the entire graph into a fixed-length feature vector of size 256 using the dgl.readout_nodes function.

#### 4.2.3 pHmin prediction module

Given the updated protein feature vectors ℎ_1_ ∈ ^1×d^ and ℎ_2_ ∈ ^1×d^, pHmin prediction module were used to generate the final predicted pH_min_.

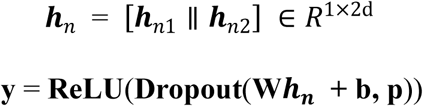

### 4.3 Performance evaluation

#### 4.3.1 Evaluating metrics

RMSE[40], PCC[41] and SPCC[42] were used to assess the performance of the model in predicting protein pHmin values. In the following formula, *y_i_* and *ȳ_i_* represent the experimental and predicted pH value of the ^th^ sample, respectively, *ŷ_i_* and 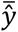 represent the average of the experimental and predicted pH values, respectively, and represents the number of samples.

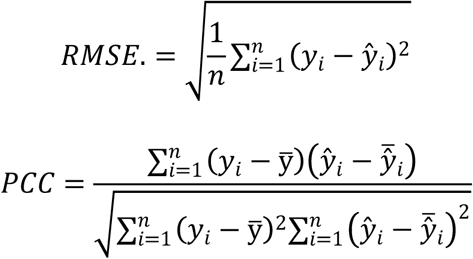

The SPCC calculation is based on the ranking of the variables, not their raw data values. This is because both *y_i_* and *ȳ_i_* need to be evaluated separately before each data point can be ranked. For identical values (i.e., identical ranks), the average rank was assigned. Therefore, for each pair of data points (*y_i_*, *ȳ_i_*), their rank difference *d_i_* was calculated: *d_i_* = rank(*y_i_*) − (*ȳ_i_*). Finally, the SPCC was calculated using the following formula:

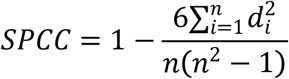

#### 4.3.2 Ablation study

To assess the impact of different components on model performance, we conducted an ablation study focusing on three key aspects: baseline graph neural network (GNN) architecture, node feature composition, and input graph types. First, to evaluate the graph convolutional network (GCN), we replaced it with two widely used GNN architectures, namely the graph attention network (GAT) [43] and the graph isomorphism network (GIN)[44]. GAT introduces an attention mechanism that adaptively assigns importance weights to neighboring nodes, while GIN aims to maximize the expressive power of GNNs to make them as powerful as the Weisfeiler-Lehman graph isomorphism test. By replacing GCN with these architectures, we aim to evaluate whether GCN provides the most effective feature extraction capability for protein acid stability prediction.

To further investigate the contribution of node features, we conducted an ablation by removing two types of sequence-derived embeddings (a 1280-dimensional feature representation extracted from the ESM-2 model[15] and a 16-dimensional physicochemical feature set, respectively) to evaluate their contribution to the accuracy of ACENet. Finally, we examined the input graph structure by selectively removing specific graph representations and retraining the model. This includes the full protein graph, which encodes backbone connectivity and structural relationships, and a surface graph constructed based on surface accessibility and geometric properties. We then evaluated whether integrating these two structural representations can provide complementary information and improve the accuracy of ACENet.

#### 4.3.3 Comparative Model Analysis

To establish a robust and rigorous benchmark for evaluating the performance of ACENet, we conducted an extensive and systematic comparative analysis of eight representative machine learning (ML) and deep learning (DL) architectures. The selected models include six classic machine learning algorithms: Random Forest (RF)[45], Extreme Gradient Boosting (XGBoost)[46], Gradient Boosting[47], Ridge Regression[48], K-Nearest Neighbors Regression (KNR)[49], and Linear Regression[50], along with two neural network architectures: Multilayer Perceptron (MLP)[51] and Convolutional Neural Network (CNN)[52]. In addition, the comparison also uses pre-trained models in the VenusFactory toolkit, including the T5 series (ProtT5-xl-uniref50)[53], BERT series (ProtBert-bfd, ProtBert-uniref50)[54], Venus-PETA series (Deep_BPE_50, Deep_BPE_400, Deep_Unigram_50, and Deep_Unigram_400)[55], and ESM series (ESM-2) [15]. All models from venusfactory (e.g., ESM-650M) used PLM to embed protein sequences, followed by pooling with a mild attention mechanism, and finally use MLP as the regression head, with the PLM module remaining frozen. This paper uses the 650M version of ESM-2. To compare with the architecture developed in this study, we also reproduced some previous methods used for pHopt prediction (EpHod[21], OphPred[19], Venus-Dream[23], opHReda[24]). This study selected architectures based on these four methods to train models from scratch to test their performance. These models were strategically selected to cover a wide range of methodological paradigms in the fields of ML and DL, from traditional ensemble methods (RF, XGBoost, Gradient Boosting), linear models (Ridge Regression, Linear Regression), distance-based approaches (KNR), to advanced neural architectures capable of capturing complex patterns in biological data (MLP, CNN), as well as state-of-the-art protein language models that have achieved remarkable success in biological big data applications.

All models included in the comparative analysis were trained and tested on the same dataset as ACENet. Hyperparameters for each model were optimized using a comprehensive grid search technique, and the optimal configuration was subsequently used to evaluate predictive performance on the test set. All models implemented using the VenusFactory toolkit were deployed with default parameters. To ensure a fair comparison, all models except the VenusFactory model and the CNN model use iFeature’s manual feature encodings from (AAC, CKSAAP)[56] and one-hot encoding for initial feature representation. The VenusFactory model can be accessed directly from https://github.com/ai4protein/VenusFactory and the required datasets were processed and could be obtained from https://huggingface.co/datasets/EzrealHu/ACENet. The code for the other models can be obtained from https://github.com/Showmake2/ACENet.

#### 4.3.4 Evaluation of the generalizability and prediction stability of ACENet

MMSeqs2[57] software and tmscoring script (https://github.com/Dapid/tmscoring) were used to calculate sequence homology (identity) and structural similarity (TM-score). The test set was divided into five sub-datasets based on the sequence identity and TM-score thresholds of 20%, 40%, 60%, and 80%, respectively, and the performance of the model was evaluated within each sub- dataset.

To investigate the impact of structural accuracy on model predictions, we collected data on monomeric proteins whose structures have been experimentally solved from the past five years of the CASP competition. Next, we used ESMFold to predict the structures of these proteins. The experimentally solved structures and their associated ESMFold predicted structures were grouped separately and fed into the model for analysis.

First, 155 microorganisms capable of growing at pH below 4 were identified from the BacDive database. Each microorganism was then manually verified and its complete proteome was retrieved from the NCBI database. Finally, only the genome sequences of 20 microorganisms are publicly available. **Table S8** lists the strain-specific growth pH ranges, optimal pH conditions, genome sequencing data, and strain identifiers. The computational workflows includes a rapid screening script to assess the availability of protein structures in the AlphaFold Protein Structure Database; when experimental structures are unavailable, we used ESMFold for structure prediction. Further, signal peptide detection was determined using SignalP 6.0, and transmembrane domain prediction was conducted using TMHMM 2.0 to determine protein subcellular localization. Proteins containing signal peptides and no transmembrane domains were designated as secreted, while the remaining proteins were classified as non-secreted. This pipeline generates a comprehensive dataset of 70,751 proteins, including source organism data, growth environment characteristics, subcellular localization predictions, and additional metadata, forming an evolutionary origin validation dataset.

### 4.4 Model interpretation

This study uses perturbation-based feature importance analysis for model interpretation. This method involves perturbing input features —such as adding noise or zeroing them out—and observing the resulting changes in model output to evaluate the contribution of each feature. To achieve residue-level interpretation, the input data is structured as a matrix ∈ ^×^, where denotes the length of the protein sequence and represents the feature dimension at each amino acid position (here, 1296). For each position in the sequence, the corresponding feature vector is perturbed by zeroing it. This creates a perturbed input, ^’^, where only the feature vector at position is modified to 0. The original input and the perturbed input ^’^ are fed into the model to obtain the original and perturbed predictions and ^’^ . The change in the prediction value, defined as Δ = − ^’^, serves as the importance score, thereby quantifying the contribution of each amino acid residue to the model’s prediction.

### 4.5 Predicting pHmin of mutant enzymes using ACENet

A comprehensive literature search was conducted using the keywords “protein engineering” and “acid stability” to find studies that presented the relative activity curves of enzymes at different pH levels. Based on relevant studies in the literature, the enzymes Amy7C, ROAmy, and CbX-CD were selected as test subjects. Amy7C is an amylase requiring enhanced acid activity in food processing and biofuel production [5]; ROAmy is a resistant starch degrading enzyme essential for the manufacture of low glycemic index foods and improves acid activity and stability by optimizing starch hydrolysis efficiency at lower pH [58]; CbX-CD is a chitosanase with potential applications in agriculture, pharmaceuticals, and food industry that must function in an acidic environment because its substrates, chitin and chitosan, are more soluble under acidic conditions [59]. We first used MMseqs2 to align the wild-type sequences of these three proteins with all proteins in the training set and calculated the sequence similarity of these three enzymes relative to the entire training data. Relative activity curves of these enzymes under different pH conditions were used to determine their pHmin values, defined as the pH at which enzyme activity first dropped below 50%. To minimize error and maximize reliability, the final pHmin values were derived from experimental data using a 0.5 pH gradient. Among these enzymes, the experimental results for Amy7C, ROAmy, and CbX-CD were chosen as the final reference for testing.

### 4.6 Implementation details

This study was implemented using PyTorch and the DGL platform. We divided the benchmark dataset into a test set with a ratio of 9:1 and used 9-fold cross validation to ensure consistency in the number of validation sets. The model was trained using the SmoothL1Loss[60] and Adam[61] optimizers, with a learning rate and weight decay set to 1e-4 and 1e-4, respectively. The model was trained for 35 epochs, with the learning rate reduced by 0.1 every 20 epochs. The batch size was set to 16. All experiments were conducted using a single NVIDIA GeForce RTX 4090 graphics card to ensure efficient computation of the benchmark dataset. Computational efficiency analysis shows that surface information extraction takes an average of 1.28 s per protein, graph generation takes 2.49 s per protein, and batch prediction (batch size=16) takes 0.74 s per batch.

## Data and code availability

The ACENet code is available at https://github.com/showmake2/ACENet. Benchmark data is sourced from https://www.brenda-enzymes.org, and the data retrieval script is from https://github.com/SysBioChalmers/DLKcat/tree/master/DeeplearningApproach/Code/preprocess. The dataset is available at https://huggingface.co/datasets/EzrealHu/ACENet/. Protein structure data are from the results of the six recent CASP (Critical Assessment of Techniques for Protein Structure Prediction) competitions.

## Acknowledgements

This work was supported by National Key R&D Program of China (2024YFA0918802), National Natural Science Foundation of China (32370101) and Beijing Natural Science Foundation (L244001).

## Author Contributions

S.H., T.P., J.S., and G.Z. contributed to conceptualizing and proposing the research idea. S.H., J.W., Z.L., and T.P. collected and curated the dataset for this work. S.H. contributed to dataset preprocessing and quality control. S.H. was responsible for designing and implementing the complete deep learning framework and conducting comprehensive performance evaluations and testing. Z.W. and T.P. provided detailed guidance and optimization for the deep learning architecture and methodology. L.Z. was responsible for constructing, training, and testing all machine learning models used in this study. S.H., Z.W., T.P., H.Y.K., and G.Z. contributed to writing and editing the manuscript. All authors reviewed the manuscript, provided critical feedback, and approved the final version for publication.

