## Supplementary material for "ACENet: a graph neural network for predicting enzyme pHmin using structure and sequence insights": Table S1-15

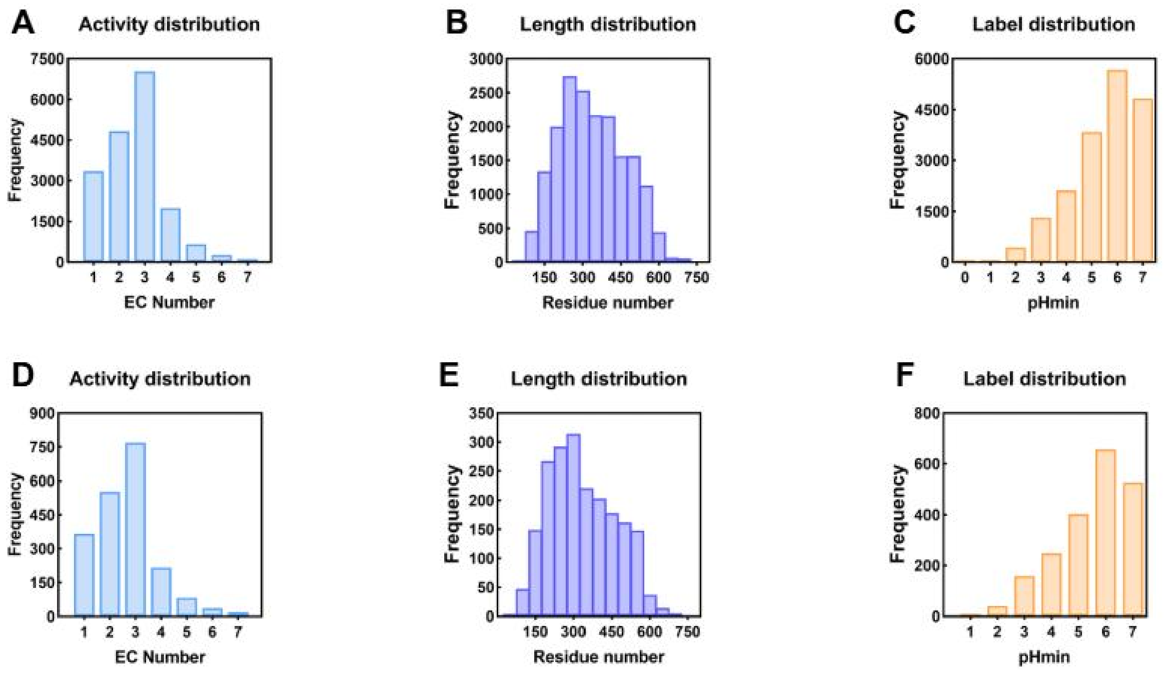


**Figure S1. The distribution of the training set(A-C) and test set(D-E).**

*EC Number(1: oxidoreductases, 2: transferases, 3: hydrolases, 4: lyases, 5: isomerases, 6: ligases, 7: translocases), sequence length and pHmin range distribution(1: pHmin≤1, 2: 1<pHmin≤2, 3: 2<pHmin≤3, 4: 3<pHmin≤4, 5: 4<pHmin≤5, 6: 5<pHmin≤6, 7: 6<pHmin≤7)

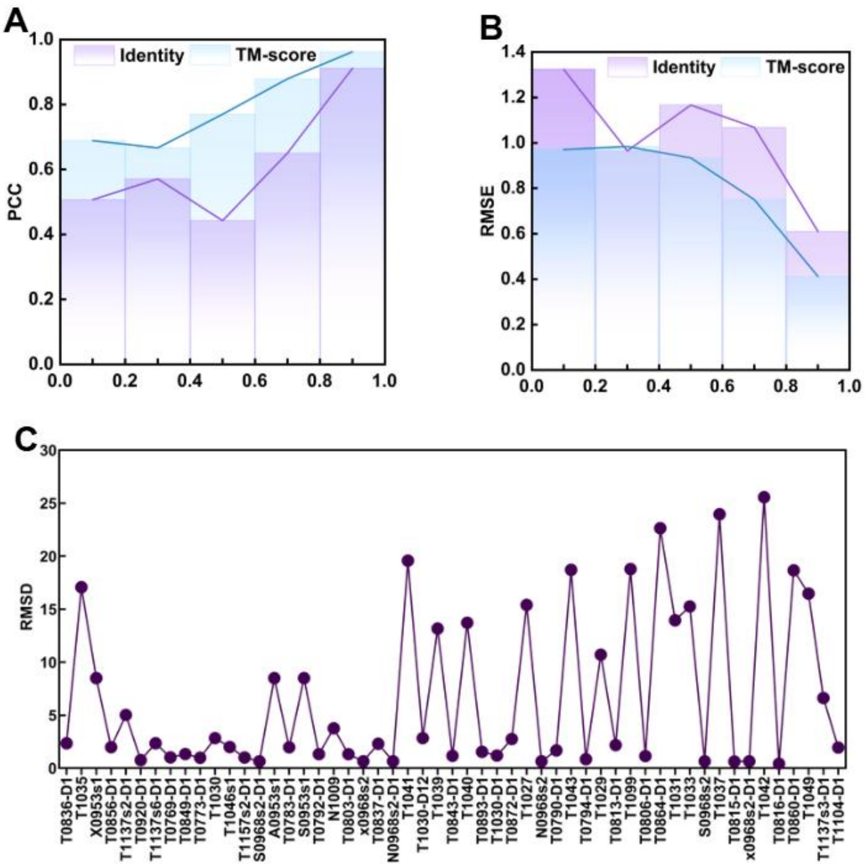


**Figure S2. Evaluation of ACENet's generalization and prediction stability. (A-B)The impact of sequence and structural similarity on model performance. (C)The Root Mean Square Deviation(RMSD) of experimentally solved structures and predicted structures.**

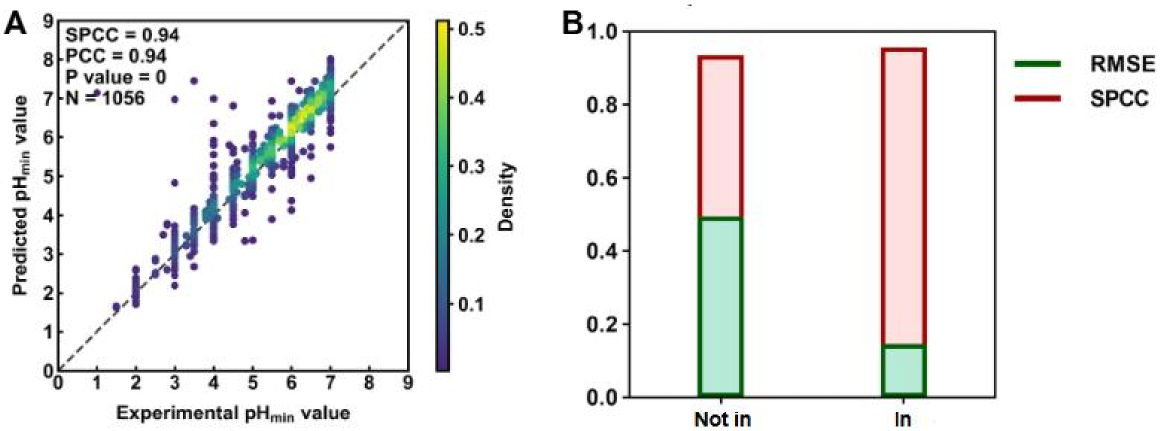


**Figure S3. Prediction of pHmin for wild-type and mutant enzymes using ACENet. (A) Prediction performance for mutant enzymes in the test set. (B) Comparison of predictive performance for test set mutants based on whether they appear in the training set or not.** "Not in" and "In" represent mutants not appearing in the training set and mutants already appearing in the training set, respectively.

**Table S1. Performance metrics of ACENet at different training epochs in the validation set**

| Epoch | SPCC | PCC | RMSE |
|---|---|---|---|
| 1 | 0.275 ± 0.040 | 0.272 ± 0.038 | 2.331 ± 0.222 |
| 3 | 0.492 ± 0.033 | 0.503 ± 0.039 | 1.576 ± 0.267 |
| 5 | 0.527 ± 0.032 | 0.532 ± 0.033 | 1.287 ± 0.117 |
| 7 | 0.602 ± 0.028 | 0.612 ± 0.026 | 1.498 ± 0.406 |
| 9 | 0.640 ± 0.064 | 0.650 ± 0.077 | 1.211 ± 0.282 |
| 11 | 0.671 ± 0.038 | 0.684 ± 0.034 | 1.333 ± 0.497 |
| 13 | 0.675 ± 0.030 | 0.687 ± 0.031 | 2.046 ± 0.930 |
| 15 | 0.708 ± 0.027 | 0.716 ± 0.026 | 1.256 ± 0.344 |
| 17 | 0.725 ± 0.034 | 0.732 ± 0.037 | 1.227 ± 0.419 |
| 19 | 0.752 ± 0.020 | 0.762 ± 0.016 | 0.956 ± 0.088 |
| 21 | 0.752 ± 0.028 | 0.760 ± 0.029 | 1.025 ± 0.145 |
| 23 | 0.760 ± 0.022 | 0.772 ± 0.022 | 0.953 ± 0.106 |
| 25 | 0.769 ± 0.016 | 0.777 ± 0.018 | 0.878 ± 0.073 |
| 27 | 0.769 ± 0.022 | 0.775 ± 0.026 | 0.978 ± 0.120 |
| 29 | 0.785 ± 0.016 | 0.795 ± 0.015 | 0.918 ± 0.172 |
| 31 | 0.787 ± 0.012 | 0.795 ± 0.009 | 0.894 ± 0.115 |
| 33 | 0.791 ± 0.011 | 0.798 ± 0.014 | 0.854 ± 0.106 |
| 35 | 0.791 ± 0.014 | 0.796 ± 0.016 | 0.827 ± 0.038 |

**Table S2. Performance metrics of ACENet at different training epochs in the test set**

| Epoch | SPCC | PCC | RMSE |
|---|---|---|---|
| 1 | 0.306 ± 0.043 | 0.303 ± 0.038 | 2.290 ± 0.233 |
| 3 | 0.523 ± 0.030 | 0.522 ± 0.043 | 1.557 ± 0.250 |
| 5 | 0.547 ± 0.026 | 0.548 ± 0.027 | 1.268 ± 0.102 |
| 7 | 0.626 ± 0.024 | 0.633 ± 0.025 | 1.477 ± 0.420 |
| 9 | 0.654 ± 0.067 | 0.663 ± 0.077 | 1.187 ± 0.274 |
| 11 | 0.689 ± 0.030 | 0.703 ± 0.030 | 1.312 ± 0.513 |
| 13 | 0.693 ± 0.027 | 0.702 ± 0.033 | 2.043 ± 0.946 |
| 15 | 0.718 ± 0.021 | 0.727 ± 0.022 | 1.237 ± 0.341 |
| 17 | 0.733 ± 0.033 | 0.743 ± 0.037 | 1.201 ± 0.416 |
| 19 | 0.756 ± 0.018 | 0.768 ± 0.016 | 0.941 ± 0.089 |
| 21 | 0.754 ± 0.022 | 0.763 ± 0.022 | 1.030 ± 0.158 |
| 23 | 0.761 ± 0.023 | 0.773 ± 0.023 | 0.960 ± 0.110 |
| 25 | 0.768 ± 0.013 | 0.779 ± 0.014 | 0.873 ± 0.062 |
| 27 | 0.769 ± 0.018 | 0.777 ± 0.019 | 0.980 ± 0.117 |
| 29 | 0.784 ± 0.008 | 0.797 ± 0.007 | 0.919 ± 0.184 |
| 31 | 0.785 ± 0.006 | 0.795 ± 0.006 | 0.890 ± 0.105 |
| 33 | 0.786 ± 0.012 | 0.798 ± 0.011 | 0.854 ± 0.099 |
| 35 | 0.787 ± 0.009 | 0.796 ± 0.009 | 0.827 ± 0.026 |

**Table S3. Ablation study of ACENet**

| Method | RMSE (test) | PCC (test) | SPCC (test) | RMSE (test<0.4) | PCC (test<0.4) | SPCC (test<0.4) |
|---|---|---|---|---|---|---|
| ACENet | 0.723 | 0.852 | 0.835 | 1.124 | 0.539 | 0.461 |
| GAT | 0.754 | 0.830 | 0.815 | 1.140 | 0.500 | 0.401 |
| GIN | 0.772 | 0.836 | 0.814 | 1.876 | 0.120 | 0.092 |
| without_ESM | 1.612 | 0.474 | 0.484 | 1.159 | 0.471 | 0.396 |
| without_phychem | 0.731 | 0.842 | 0.827 | 1.236 | 0.446 | 0.358 |
| without_full_graph | 0.786 | 0.836 | 0.823 | 5.839 | 0.455 | 0.348 |
| without_surface_graph | 0.733 | 0.840 | 0.824 | 1.139 | 0.499 | 0.427 |

**Table S4. Performance comparison of the best model from each method**

| Model | Feature | test | SPCC | |
|---|---|---|---|---|
| Venus-Dream | | Test | 0.7469 | pHopt model |
| | | Test<40% | 0.3874 | |
| OpHReda | | Test | 0.6543 | |
| | | Test<40% | 0.3539 | |
| OphPred | | Test | 0.5972 | |
| | | Test<40% | 0.2706 | |
| Ephod | | Test | 0.7978 | |
| | | Test<40% | 0.3992 | |
| XGBoost | iFeature | Test | 0.7318 | Machine learning |
| | | Test<40% | 0.17 | |
| XGBoost | One-hot | Test | 0.6761 | |
| | | Test<40% | 0.03 | |
| RandomForest | iFeature | Test | 0.779 | |
| | | Test<40% | 0.37 | |
| RandomForest | One-hot | Test | 0.7151 | |
| | | Test<40% | 0.11 | |
| GradientBoosting | iFeature | Test | 0.5907 | |
| | | Test<40% | 0.33 | |
| GradientBoosting | One-hot | Test | 0.5451 | |
| | | Test<40% | 0.15 | |
| Ridge | iFeature | Test | 0.41 | |
| | | Test<40% | 0.39 | |
| Ridge | One-hot | Test | 0.5243 | |
| | | Test<40% | 0.1 | |
| KNR | iFeature | Test | 0.7774 | |
| | | Test<40% | 0.27 | |
| KNR | One-hot | Test | 0.672 | |
| | | Test<40% | 0.2 | |
| LinearRegression | iFeature | Test | 0.4178 | |
| | | Test<40% | 0.41 | |
| LinearRegression | One-hot | Test | 0.4248 | |
| | | Test<40% | 0.19 | |
| MLP | iFeature | Test | 0.6218 | Deep learning |
| | | Test<40% | 0.27 | |
| MLP | One-hot | Test | 0.6981 | |
| | | Test<40% | 0.31 | |
| CNN | One-hot | Test | 0.7678 | |
| | | Test<40% | 0.33 | |

| | | | | |
|---|---|---|---|---|
| Light attention(freeze) | Ankh-base | Test | 0.6993 | PLMs |
| | | Test<40% | 0.2755 | |
| | Deep_BPE_50 | Test | 0.7751 | |
| | | Test<40% | 0.1706 | |
| | Deep_BPE_400 | Test | 0.7785 | |
| | | Test<40% | 0.4198 | |
| | Deep_Unigram_50 | Test | 0.6721 | |
| | | Test<40% | 0.2322 | |
| | Deep_Unigram_400 | Test | 0.7794 | |
| | | Test<40% | 0.2833 | |
| | ESM-2 650M | Test | 0.7931 | |
| | | Test<40% | 0.4337 | |
| | ProtBert-bfd | Test | 0.7438 | |
| | | Test<40% | 0.4043 | |
| | ProtBert-uniref50 | Test | 0.7056 | |
| | | Test<40% | 0.3055 | |
| | ProtT5-xl-uniref50 | Test | 0.6986 | |
| | | Test<40% | 0.3853 | |
| | ACENet | Test | 0.8348 | |
| | | Test<40% | 0.4649 | |

**Table S5. Evaluation of the generalizability of the model to the test set at different sequence similarity levels**

| Identity | RMSE | PCC | SPCC | P-value |
|---|---|---|---|---|
| 0.0-0.2 | 0.971 | 0.689 | 0.639 | 1.620E-45 |
| 0.2-0.4 | 0.984 | 0.666 | 0.654 | 1.620E-50 |
| 0.4-0.6 | 0.934 | 0.770 | 0.766 | 1.710E-31 |
| 0.6-0.8 | 0.750 | 0.879 | 0.861 | 2.340E-51 |
| 0.8-1.0 | 0.413 | 0.962 | 0.950 | 0 |

**Table S6. Evaluation of the generalizability of the model at different levels of structural similarity (TM-score) using the test set**

| TM-score | RMSE | PCC | SPCC | P-value |
|---|---|---|---|---|
| 0.0-0.2 | 1.324 | 0.507 | 0.349 | 3.069E-05 |
| 0.2-0.4 | 0.965 | 0.571 | 0.545 | 4.487E-10 |
| 0.4-0.6 | 1.167 | 0.443 | 0.418 | 2.991E+00 |
| 0.6-0.8 | 1.069 | 0.650 | 0.664 | 2.735E-18 |
| 0.8-1.0 | 0.611 | 0.912 | 0.899 | 0 |

**Table S7. Performance evaluation of ACENet across different EC Numbers in the training and test set**

| EC number | RMSE(Test) | RMSE(Train) |
|---|---|---|
| 1 | 0.7854 | 0.4846 |
| 2 | 0.6898 | 0.4297 |
| 3 | 0.7183 | 0.4613 |
| 4 | 0.7545 | 0.4659 |
| 5 | 0.7189 | 0.4655 |
| 6 | 0.5068 | 0.3655 |
| 7 | 0.5406 | 0.4647 |
| All | 0.7227 | 0.4570 |

**Table S8. Performance evaluation of ACENet in different pH stability ranges in the training set**

| pH | RMSE | Num |
|---|---|---|
| 0<pHmin<=3 | 0.586 | 1694 |
| 3<pHmin<=4 | 0.431 | 2110 |
| 4<pHmin<=5 | 0.364 | 3429 |
| 5<pHmin<=6 | 0.357 | 5731 |
| 6<pHmin<=7 | 0.438 | 5282 |

**Table S9. Performance evaluation of ACENet in different pH stability ranges in the test set**

| pH | RMSE | Num |
|---|---|---|
| 0<pHmin<=3 | 1.257 | 192 |
| 3<pHmin<=4 | 0.871 | 248 |
| 4<pHmin<=5 | 0.628 | 360 |
| 5<pHmin<=6 | 0.553 | 653 |
| 6<pHmin<=7 | 0.629 | 585 |

**Table S10. Prediction of pHmin in secreted and non-secreted proteins of Gram-positive and Gram-negative bacteria**

| ID | Strain_number_header | Species | Growth pH | Optimum pH | Num_proteins |
|---|---|---|---|---|---|
| 164491 | HSLZ-75(=JCM 33270) | *Acetilactobacillus jinshanensis* | 3.0-5.0 | 4 | 1667 |
| 166226 | G45-3 | *Acidibrevibacterium fodinaquatile* | 2.5-5 | 4 | 3617 |
| 131 | ATCC 51196 | *Acidobacterium capsulatum* | 3.0-6.0 | None | 4380 |
| 23706 | CBMB205 | *Bacillus velezensis* | 2.0-10.0 | 7-7.5 | 3895 |
| 2090 | DSM 15908 | *Caldisphaera lagunensis* | 2.3-5.4 | 3.5-4.0 | 1807 |
| 141129 | S5(T) (=JCM 30642; =VKM B-2941) | *Cuniculiplasma divulgatum* | 0.5-4.0 | 1.0-1.2 | 2248 |
| 158304 | DSM 104462 | *Tunturiibacter lichenicola* | 3.4-7.0 | 4.3-5.6 | 4695 |
| 17689 | YUAN-3 | *Ethanoligenens harbinense* | 3.5-9 | 4.5-5.0 | 3343 |
| 22901 | MP5ACTX9 | *Granulicella tundricola* | 3.5-6.5 | 5 | 5444 |
| 6089 | c2 | *Halothiobacillus neapolitanus* | 3-8.5 | 6.5-6.9 | 2622 |
| 4189 | DSM 13165 | *Ignicoccus islandicus* | 3.8-6.5 | 5.8 | 1746 |
| 141151 | WiKim39 | *Companilactobacillus allii* | 4.5-9.0 | None | 2709 |
| 166412 | ZY170218 | *Mannheimia pernigra* | 3.0-11.0 | 9 | 2115 |
| 1673 | BW863 | *Methylovirgula ligni* | 3.1-6.5 | 4.5-5.0 | 3804 |
| 132351 | DSM 25168 | *Occallatibacter riparius* | 3.5-8.5 | 5.0-6.5 | 5685 |
| 132417 | PX4 | *Paludisphaera borealis* | 3.5-6.5 | 5-5.5 | 6544 |
| 158708 | BN5 | *Paraburkholderia aromaticivorans* | 3.0-7.0 | 5.0-6.0 | 8527 |
| 158656 | HS-3(=JCM 32116) | *Saccharolobus caldissimus* | 1.5-6 | 3 | 3728 |
| 140590 | HS-1 | *Sulfodiicoccus acidiphilus* | 1.4-5.5 | 3.0-3.5 | 3077 |
| 140262 | 3127-1 | *Thermodesulfobium acidiphilum* | 3.7-6.5 | 4.8-5.0 | 1947 |

**Table S11. Residue importance analysis in glutathione transferase and carbonic anhydrase**

| Amino Acid | Carbonic anhydrase | | | | | Glutathione transferase | | | | | |
|---|---|---|---|---|---|---|---|---|---|---|---|
| | pH 4.0 | pH 4.5 | pH 5.0 | pH 6.0 | Mean | pH 4.0 | pH 5.0 | pH 6.0 | pH 6.5 | pH 7.0 | Mean |
| A | 0.020 | 0.080 | 0.040 | 0.180 | 0.080 | 0.020 | 0.100 | 0.100 | 0.080 | 0.100 | 0.080 |
| V | 0.020 | 0.040 | 0.040 | 0.000 | 0.025 | 0.000 | 0.060 | 0.060 | 0.040 | 0.080 | 0.048 |
| L | 0.060 | 0.060 | 0.080 | 0.140 | 0.085 | 0.060 | 0.080 | 0.060 | 0.060 | 0.020 | 0.056 |
| I | 0.020 | 0.040 | 0.000 | 0.000 | 0.015 | 0.080 | 0.020 | 0.020 | 0.020 | 0.040 | 0.036 |
| M | 0.060 | 0.060 | 0.040 | 0.020 | 0.045 | 0.100 | 0.040 | 0.040 | 0.040 | 0.040 | 0.052 |
| F | 0.000 | 0.060 | 0.040 | 0.040 | 0.035 | 0.040 | 0.020 | 0.080 | 0.100 | 0.040 | 0.056 |
| Y | 0.000 | 0.040 | 0.040 | 0.020 | 0.025 | 0.100 | 0.100 | 0.040 | 0.000 | 0.040 | 0.056 |
| W | 0.000 | 0.020 | 0.000 | 0.020 | 0.010 | 0.020 | 0.000 | 0.020 | 0.020 | 0.020 | 0.016 |
| S | 0.040 | 0.180 | 0.140 | 0.140 | 0.125 | 0.000 | 0.120 | 0.040 | 0.060 | 0.160 | 0.076 |
| T | 0.000 | 0.060 | 0.180 | 0.100 | 0.085 | 0.020 | 0.000 | 0.020 | 0.020 | 0.120 | 0.036 |
| N | 0.000 | 0.000 | 0.080 | 0.000 | 0.020 | 0.080 | 0.040 | 0.040 | 0.060 | 0.020 | 0.048 |
| Q | 0.020 | 0.060 | 0.020 | 0.060 | 0.040 | 0.040 | 0.080 | 0.020 | 0.060 | 0.040 | 0.048 |
| G | 0.160 | 0.040 | 0.060 | 0.120 | 0.095 | 0.060 | 0.040 | 0.080 | 0.040 | 0.000 | 0.044 |
| P | 0.020 | 0.000 | 0.060 | 0.000 | 0.020 | 0.060 | 0.040 | 0.040 | 0.080 | 0.040 | 0.052 |
| C | 0.020 | 0.020 | 0.040 | 0.040 | 0.030 | 0.000 | 0.020 | 0.000 | 0.000 | 0.000 | 0.004 |
| D | 0.100 | 0.020 | 0.040 | 0.020 | 0.045 | 0.100 | 0.000 | 0.060 | 0.060 | 0.100 | 0.064 |
| E | 0.260 | 0.100 | 0.040 | 0.020 | 0.105 | 0.120 | 0.040 | 0.140 | 0.120 | 0.040 | 0.092 |
| H | 0.060 | 0.040 | 0.000 | 0.040 | 0.035 | 0.020 | 0.020 | 0.000 | 0.000 | 0.000 | 0.008 |
| K | 0.100 | 0.040 | 0.040 | 0.000 | 0.045 | 0.000 | 0.100 | 0.040 | 0.040 | 0.080 | 0.052 |
| R | 0.040 | 0.040 | 0.020 | 0.040 | 0.035 | 0.080 | 0.080 | 0.100 | 0.100 | 0.020 | 0.076 |

**Table S12. Amino acid frequencies in glutathione transferase and carbonic anhydrase**

| Amino Acid | Carbonic anhydrase | | | | | Glutathione transferase | | | | | |
|---|---|---|---|---|---|---|---|---|---|---|---|
| | pH 4.0 | pH 4.5 | pH 5.0 | pH 6.0 | Mean | pH 4.0 | pH 5.0 | pH 6.0 | pH 6.5 | pH 7.0 | Mean |
| A | 0.054 | 0.087 | 0.068 | 0.077 | 0.071 | 0.064 | 0.046 | 0.070 | 0.071 | 0.074 | 0.065 |
| V | 0.008 | 0.063 | 0.079 | 0.047 | 0.050 | 0.041 | 0.050 | 0.070 | 0.060 | 0.083 | 0.061 |
| L | 0.041 | 0.102 | 0.091 | 0.145 | 0.095 | 0.106 | 0.100 | 0.126 | 0.159 | 0.134 | 0.125 |
| I | 0.095 | 0.032 | 0.028 | 0.015 | 0.042 | 0.069 | 0.104 | 0.061 | 0.044 | 0.042 | 0.064 |
| M | 0.041 | 0.039 | 0.020 | 0.015 | 0.029 | 0.046 | 0.025 | 0.047 | 0.044 | 0.032 | 0.039 |
| F | 0.086 | 0.039 | 0.040 | 0.030 | 0.049 | 0.046 | 0.033 | 0.028 | 0.033 | 0.023 | 0.033 |
| Y | 0.049 | 0.024 | 0.032 | 0.015 | 0.030 | 0.060 | 0.046 | 0.042 | 0.044 | 0.046 | 0.048 |
| W | 0.070 | 0.024 | 0.016 | 0.012 | 0.030 | 0.014 | 0.008 | 0.005 | 0.006 | 0.009 | 0.008 |
| S | 0.103 | 0.126 | 0.060 | 0.100 | 0.097 | 0.028 | 0.062 | 0.051 | 0.017 | 0.074 | 0.046 |
| T | 0.062 | 0.063 | 0.087 | 0.047 | 0.065 | 0.073 | 0.042 | 0.033 | 0.044 | 0.079 | 0.054 |
| N | 0.037 | 0.047 | 0.071 | 0.018 | 0.043 | 0.055 | 0.050 | 0.028 | 0.028 | 0.019 | 0.036 |
| Q | 0.041 | 0.032 | 0.036 | 0.050 | 0.040 | 0.028 | 0.025 | 0.028 | 0.033 | 0.023 | 0.027 |
| G | 0.054 | 0.047 | 0.068 | 0.094 | 0.066 | 0.051 | 0.046 | 0.056 | 0.055 | 0.037 | 0.049 |
| P | 0.021 | 0.071 | 0.071 | 0.086 | 0.062 | 0.064 | 0.071 | 0.047 | 0.044 | 0.056 | 0.056 |
| C | 0.037 | 0.008 | 0.012 | 0.009 | 0.016 | 0.005 | 0.029 | 0.009 | 0.011 | 0.000 | 0.011 |
| D | 0.054 | 0.032 | 0.024 | 0.041 | 0.038 | 0.073 | 0.054 | 0.056 | 0.071 | 0.083 | 0.068 |
| E | 0.025 | 0.039 | 0.056 | 0.103 | 0.056 | 0.060 | 0.071 | 0.070 | 0.082 | 0.037 | 0.064 |
| H | 0.070 | 0.032 | 0.040 | 0.050 | 0.048 | 0.018 | 0.029 | 0.023 | 0.017 | 0.019 | 0.021 |
| K | 0.017 | 0.055 | 0.056 | 0.009 | 0.034 | 0.055 | 0.066 | 0.102 | 0.082 | 0.111 | 0.084 |
| R | 0.037 | 0.039 | 0.048 | 0.038 | 0.041 | 0.046 | 0.046 | 0.051 | 0.055 | 0.019 | 0.043 |

**Table S13. Residue importance analysis in the entire dataset**

| Amino Acid | pHmin | | | | | Mean |
|---|---|---|---|---|---|---|
| | <3 | 3 | 4 | 5 | 6 | |
| A | 0.108 | 0.109 | 0.107 | 0.104 | 0.115 | 0.109 |
| C | 0.024 | 0.028 | 0.025 | 0.022 | 0.020 | 0.022 |
| D | 0.058 | 0.065 | 0.073 | 0.076 | 0.072 | 0.072 |
| E | 0.054 | 0.062 | 0.077 | 0.087 | 0.094 | 0.084 |
| F | 0.039 | 0.036 | 0.034 | 0.033 | 0.030 | 0.033 |
| G | 0.067 | 0.059 | 0.070 | 0.065 | 0.066 | 0.066 |
| H | 0.020 | 0.024 | 0.026 | 0.026 | 0.029 | 0.027 |
| I | 0.047 | 0.045 | 0.039 | 0.043 | 0.039 | 0.041 |
| K | 0.039 | 0.050 | 0.051 | 0.057 | 0.059 | 0.055 |
| L | 0.081 | 0.078 | 0.058 | 0.052 | 0.052 | 0.057 |
| M | 0.039 | 0.040 | 0.042 | 0.050 | 0.046 | 0.046 |
| N | 0.053 | 0.044 | 0.040 | 0.033 | 0.031 | 0.036 |
| P | 0.038 | 0.035 | 0.034 | 0.031 | 0.028 | 0.030 |
| Q | 0.050 | 0.054 | 0.047 | 0.055 | 0.057 | 0.055 |
| R | 0.032 | 0.034 | 0.044 | 0.049 | 0.056 | 0.050 |
| S | 0.098 | 0.095 | 0.095 | 0.087 | 0.087 | 0.091 |
| T | 0.058 | 0.051 | 0.049 | 0.047 | 0.040 | 0.045 |
| V | 0.042 | 0.042 | 0.041 | 0.041 | 0.041 | 0.042 |
| W | 0.014 | 0.015 | 0.019 | 0.016 | 0.015 | 0.016 |
| Y | 0.040 | 0.034 | 0.030 | 0.027 | 0.023 | 0.027 |

**Table S14. Amino acid frequencies in the entire dataset**

| Amino Acid | pHmin | | | | | Mean |
|---|---|---|---|---|---|---|
| | <3 | 3 | 4 | 5 | 6 | |
| A | 0.077 | 0.084 | 0.086 | 0.088 | 0.092 | 0.089 |
| C | 0.016 | 0.015 | 0.015 | 0.014 | 0.014 | 0.014 |
| D | 0.058 | 0.060 | 0.057 | 0.057 | 0.056 | 0.056 |
| E | 0.046 | 0.050 | 0.055 | 0.061 | 0.065 | 0.060 |
| F | 0.045 | 0.043 | 0.041 | 0.041 | 0.039 | 0.040 |
| G | 0.087 | 0.081 | 0.083 | 0.079 | 0.078 | 0.080 |
| H | 0.022 | 0.024 | 0.024 | 0.025 | 0.026 | 0.025 |
| I | 0.053 | 0.050 | 0.053 | 0.057 | 0.057 | 0.056 |
| K | 0.042 | 0.048 | 0.050 | 0.052 | 0.052 | 0.051 |
| L | 0.082 | 0.088 | 0.085 | 0.093 | 0.097 | 0.092 |
| M | 0.021 | 0.021 | 0.022 | 0.026 | 0.026 | 0.024 |
| N | 0.049 | 0.052 | 0.048 | 0.042 | 0.038 | 0.042 |
| P | 0.054 | 0.051 | 0.049 | 0.048 | 0.048 | 0.049 |
| Q | 0.039 | 0.037 | 0.036 | 0.037 | 0.037 | 0.037 |
| R | 0.036 | 0.043 | 0.047 | 0.049 | 0.053 | 0.049 |
| S | 0.080 | 0.071 | 0.068 | 0.059 | 0.057 | 0.062 |
| T | 0.068 | 0.058 | 0.058 | 0.055 | 0.053 | 0.055 |
| V | 0.068 | 0.066 | 0.067 | 0.072 | 0.071 | 0.070 |
| W | 0.018 | 0.019 | 0.018 | 0.014 | 0.013 | 0.015 |
| Y | 0.042 | 0.039 | 0.040 | 0.034 | 0.031 | 0.034 |

**Table S15. Predicted pHmin values of three enzymes and their corresponding mutants**

| Type | Enzyme | pHmin_exper | rank_exper | pHmin_pred | rank_pred | SPCC↑ | | RMSE↓ | |
|---|---|---|---|---|---|---|---|---|---|
| Amy7C | WT | 3.5 | 4 | 2.770 | 4 | 1.00 | 0.52 (mean) | 0.82 | 0.66 (mean) |
| | R172K | 4 | 6 | 2.799 | 6 | | | | |
| | A270K | 3.5 | 3 | 2.771 | 3 | | | | |
| | N271H | 3 | 1 | 2.757 | 1 | | | | |
| | A270K; N271H | 3 | 2 | 2.765 | 2 | | | | |
| | R172K; N271H | 4 | 5 | 2.797 | 5 | | | | |
| ROAmy | WT | 4 | 7 | 2.997 | 6 | 0.75 | | 0.74 | |
| | K71E | 4 | 6 | 2.988 | 5 | | | | |
| | G136D | 3.5 | 2 | 2.980 | 3 | | | | |
| | A144Y | 3.5 | 3 | 2.960 | 2 | | | | |
| | V174R | 3 | 1 | 3.000 | 1 | | | | |
| | T253E | 3.5 | 4 | 3.010 | 7 | | | | |
| | I276P | 4 | 5 | 2.990 | 4 | | | | |
| CbX-CD | WT | 4.5 | 10 | 3.700 | 3 | 0.15 | | 0.45 | |
| | S56D | 4.5 | 8 | 3.876 | 8 | | | | |
| | A166E | 4 | 7 | 3.796 | 7 | | | | |
| | D176Y | 4.5 | 9 | 3.748 | 4 | | | | |
| | Q177E | 4 | 6 | 3.760 | 6 | | | | |
| | A166E; Q177E | 3.5 | 2 | 3.778 | 9 | | | | |
| | S56D; Q177E | 4 | 4 | 3.676 | 2 | | | | |
| | S56D; A166E; Q177E | 3.5 | 3 | 3.780 | 5 | | | | |
| | S56D; A166E | 4 | 5 | 3.725 | 3 | | | | |
| | S56D; A166E; D176Y; Q177E | 3.5 | 1 | 3.635 | 1 | | | | |
